# Extracellular disposal of nuclear waste by APP: a protective mechanism impaired in Alzheimer’s disease

**DOI:** 10.1101/2024.02.10.579739

**Authors:** Godfried Dougnon, Takayoshi Otsuka, Yuka Nakamura, Akiko Sakai, Tomoyuki Yamanaka, Noriko Matsui, Asa Nakahara, Ai Ito, Atsushi Hatano, Masaki Matsumoto, Hironaka Igarashi, Akiyoshi Kakita, Masaki Ueno, Hideaki Matsui

## Abstract

Although the amyloid beta (Aβ) hypothesis^1^ has long been central to Alzheimer’s disease (AD) research, effective therapeutic strategies remain elusive^2,3^. Here we re-evaluate the functions of amyloid precursor protein (APP) and reveal its critical function in protecting against nuclear impairment-induced cell death and inflammation^4,5^. Overexpression of APP mitigated etoposide or lamin A knockdown-induced nuclear damage, while APP removal or mutations exacerbated these effects. Interestingly, neurons differentiated from induced pluripotent stem cells (iPSCs) exhibited similar patterns, and notably, familial AD-associated mutant APP failed to confer protection against nuclear impairment. We identify APP’s interaction with a cytoplasmic structure of nuclear origin, termed “nuclear waste”, and propose its role in extracellular waste disposal. Intriguingly, cells lacking APP showed impaired nuclear waste clearance, leading to abnormal cytoplasmic accumulation of the nuclear waste. Similarly, neuron-specific APP overexpression using adeno-associated virus (AAV) in mice reduced neuronal death and inflammation caused by nuclear damage. Conversely, shRNA-mediated APP exacerbated these effects, and mutant APP associated with familial AD lacked protective effects. Moreover, postmortem analysis of AD brains revealed accumulation of abnormal nuclear waste in the neurocytoplasm, irregular nuclear morphology, and reduced APP levels per neuron. Our data underscore APP’s crucial role in disposing of nuclear waste, maintaining cellular homeostasis, and suggest its dysregulation as a potential contributor to AD pathogenesis. Restoring APP waste clearance in AD could be a promising target for disease-modifying therapies.

## Introduction

The nucleus, a robust fortress of genetic information and regulatory processes, has the remarkable role of controlling cell fate and function^6^. Yet, notwithstanding its strong architecture and defense mechanisms, the principal cell element is not impervious to perturbations, and disruptions to its integrity can unleash a cascade of events that ultimately compromise cellular well-being. Nuclear impairment can be induced by various factors such as aging, chemical treatment or genetic manipulation^4,5,7^, and such stressors have the capability of inflicting significant damage upon the cell, through a series of cellular responses, including inflammation and cellular senescence. These consequences bear relevance to neurodegenerative conditions such as Alzheimer’s disease (AD), where cellular health and integrity are progressively and drastically compromised. Amyloid precursor protein (APP), a protein nowadays recognized for its prominence in AD due to its cleavage to form amyloid beta (Aβ) peptides-the amyloid hypothesis^1^-, has attracted much attention. Indeed, the pathology of AD is renowned for the deposition of Aβ plaques and neurofibrillary tangles in the neocortex and the hippocampus, and it is commonly recognized that accumulation of Aβ-ultimate product of APP-engenders a cascade of harmful events leading to neuronal inflammation and cell death^8^. Nonetheless, despite extensive research toward the amyloidogenic theory of AD, effective therapies against AD remain a subject of intense scrutiny and inquiry. There is clear evidence that little did we know about the intriguing APP and its functions in neurodegenerative illnesses^9–11^.

Here, we undertook a multipronged approach to provide a body of evidence that APP could be a linchpin in cellular defenses against nuclear impairment and neurodegeneration.

Encompassing in vitro cellular studies, experiments using human neurons differentiated from induced pluripotent stem cells (iPSCs), in vivo investigations using mice, and thoroughly analyzing human postmortem samples, we unravel the intricate mechanisms whereby APP safeguards cellular integrity in the face of nuclear insults and proceed to nuclear waste clearance out of the cell.

## Results

### Wild type APP but not AD-related mutant protects cells from senescence-associated nuclear damage

To investigate the role of APP in cellular response to senescence-associated nuclear damage, we first conducted experiments using Hela cells. We utilized a plasmid expressing APP695, which is predominantly produced within neurons in the brain^12,13^. First, lamin A (LMNA) was knocked down using siRNA which induces nuclear membrane fragility as seen in senescence^5,7^. Knockdown of LMNA resulted in decreased cell survival, whereas overexpression of wild type APP (APP WT) rescued it (Fig. 1a). Overexpressing mutant APP associated with AD (hereafter, APP Swe/Ind) did not exhibit such a protective effect and on the contrary, led to a decrease in cell survival (Fig. 1a). Recent studies have reported that nuclear damage can cause leakage of nuclear contents into the cytoplasm, triggering inflammation and cellular senescence^14,15^. In this context, inflammatory molecules, such as IL6 and IFNβ, significantly decreased upon APP WT overexpression; however, such reduction was smaller in APP Swe/Ind overexpression (Fig. 1b). Additionally, western blot analysis revealed that γH2AX, a well-established marker of DNA double strand breaks (DSBs)^16^, increased upon LMNA reduction; nevertheless, γH2AX levels were significantly reduced by APP WT overexpression, suggesting that nuclear damage was mitigated (Fig. 1c). Next, we asked if APP overexpression would prevent cell death subsequent to DNA damage. We investigated different markers of cell death, including cleaved PARP1, cleaved Gasdermin D and cleaved caspase-3, which all levels increased upon LMNA reduction but drastically decreased when APP WT was overexpressed, indicating a strong inhibition of cell death (Fig. 1c). Conversely, no such protective effects were observed with APP Swe/Ind (Fig. 1c).

**Fig. 1.**
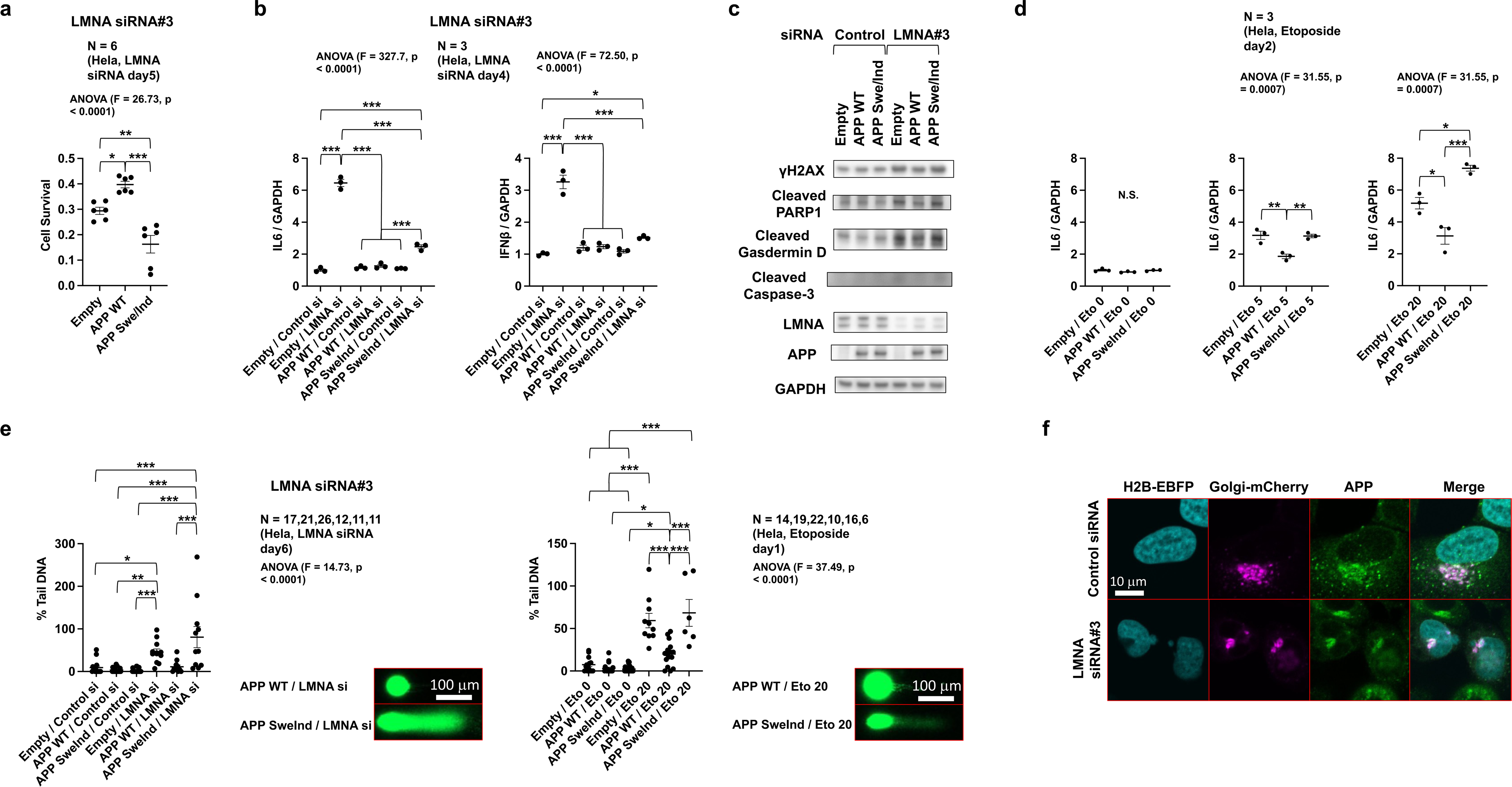
Wild type APP confers cellular protection against nuclear damage. **a**, APP WT exhibits cellular protection against LMNA siRNA-mediated toxicity. The graph shows cell viability, with values indicating the ratio relative to control siRNA (day 5). **b**, Observed intracellular inflammatory response assessed for IL6 and IFNβ following LMNA siRNA treatment, using RT-qPCR. Values represent the data averaged to empty plasmid - control siRNA (day 4). Statistical analysis: ordinary one-way ANOVA with Bonferroni posthoc test for multiple comparisons. **c**, Western blotting analysis of DNA damage and/or cell death markers including γH2AX, cleaved caspase-3, and cleaved Gasdermin D under LMNA siRNA condition (day 4). Uncropped immunoblots are provided in Supplemental Fig. 1. Reduction of LMNA by siRNA leads to an increase in several molecules related to DNA damage and cell death, which are suppressed by APP WT. However, this effect is lost in the mutant APP associated with AD (APP Swe/Ind). **d**, Intracellular inflammatory response assessed for IL6 following nuclear damage caused by etoposide (0, 5, or 20 μM, day 2), using RT-qPCR. Values represent the data averaged to the untreated empty plasmid. APP WT exhibits cellular protection against etoposide toxicity, whereas this effect is absent in APP Swe/Ind. Statistical analysis: ordinary one-way ANOVA with Bonferroni posthoc test for multiple comparisons. **e**, Comet assay results for LMNA siRNA or etoposide-treated cells. Left, LMNA siRNA (day 6). Right, etoposide treatment (0 or 20 μM, day 1). APP WT suppresses DNA damage in both cases, whereas this protective effect is lost in APP Swe/Ind. Statistical analysis: ordinary one-way ANOVA with Bonferroni posthoc test for multiple comparisons. **f**, Subcellular localization of APP during LMNA siRNA treatment (day 3). APP predominantly localizes to the Golgi apparatus. Upon LMNA siRNA-mediated nuclear envelope fragility, APP accumulates at sites presumed to be vulnerable regions of the nuclear envelope. Data are mean ± s.e.m. The exact *P*-values of comparison are presented in the figures. N.S. non-significant, \**P* < 0.05, \*\**P* < 0.01 and \*\*\**P* < 0.001.

To corroborate our findings that APP exhibits protective effects upon nuclear damage, we next treated Hela cells with etoposide, a widely used chemotherapy drug that induces DNA DSBs by multiple mechanisms and shows a generalized toxicity profile including neurologic damage and inflammatory responses^17–19^. Consistent with our previous findings, APP WT overexpression reduced the inflammatory response triggered by etoposide treatment (Fig. 1d). Notably, APP Swe/Ind demonstrated no protective effect against etoposide-induced nuclear damage (Fig. 1d). We further confirmed nuclear damage using the comet assay, and as expected, LMNA reduction or etoposide treatment led to a significant nuclear damage. Again, APP WT overexpression reduced the DNA damage, whereas this protective effect was absent with APP Swe/Ind (Fig. 1e). Next, we investigated the subcellular localization of APP under conditions of nuclear damage using immunostaining, and results demonstrated that APP accumulated around the deformed nucleus, alongside with the Golgi apparatus (Fig. 1f). This indicated that APP might have some important functions dealing with nuclear damage.

In summary, our findings demonstrate that APP exerts a protective role against nuclear damage; however, this protective function is impaired in the mutant APP Swe/Ind.

### Loss of APP function causes cellular vulnerability to senescence-associated nuclear damage

Next, using CRISPR-Cas9 technology, we established three Hela cell lines harboring APP genome mutations. Two lines (316-34 and 316-44) were knockout (KO) mutants with no detectable protein expression, and the third cell line (316-15) had a mutation with no frameshift in APP, in which most amino acids of the APP protein were translated (Fig. 2a,b and Extended Data Fig. 1a). In this third line (316-15), the wild type localization of APP to the Golgi apparatus was lost, and the mutant lines showed a diffuse localization of APP in the cytoplasm, resulting in lack of Aβ production (Fig. 2a,b). Interestingly, all mutants exhibited a significant decrease in cell survival following etoposide treatment or LMNA reduction (Fig. 2c). Importantly, western blot analysis of cell death markers revealed a pronounced increase in cleaved Gasdermin D and cleaved caspase-3 levels in all mutants compared to the WT cells after etoposide treatment or LMNA reduction (Fig. 2d). Similarly, inflammatory-related molecules, such as IL6 and IL1β, showed increased expression in the mutants compared to the WT cells following etoposide treatment or LMNA reduction (Fig. 2e). Anew, the abnormalities observed in the APP KO cells (316-34) were rescued by APP WT overexpression but not by mutant APP Swe/Ind (Fig. 2f). RNA sequencing analysis of the etoposide-treated APP mutant cells revealed primarily dysregulation in pathways related to DNA quality control and/or extracellular secretion (Extended Data Fig. 1b,c).

**Fig. 2.**
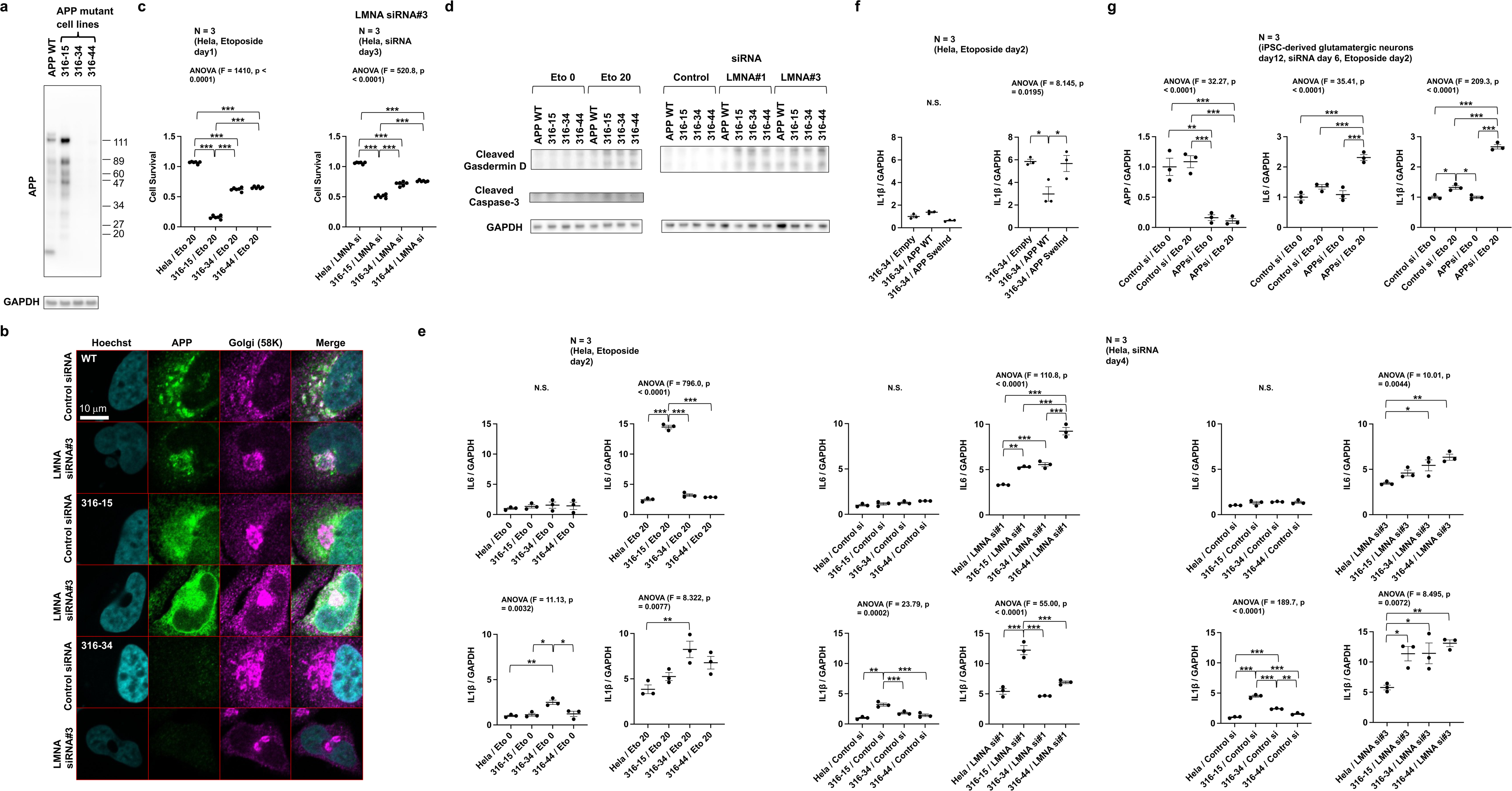
Loss of endogenous APP abolishes cellular protection against nuclear damage. **a**, Western blot analysis of APP mutant cell lines (316-15, 316-34, and 316-44) using CRISPR-Cas9 and Hela cell. 316-34 and 316-44 completely lack APP expression, while 316-15 harbors an in-frame mutation, retaining most amino acids of APP but without Aβ secretion. Uncropped immunoblot is provided in Supplemental Fig. 2a. **b**, Subcellular localization of endogenous APP during LMNA siRNA treatment (day 3). Endogenous APP predominantly localizes to the Golgi apparatus, and upon LMNA siRNA-mediated nuclear fragility, it accumulates at vulnerable regions of the nuclear envelope. In 316-34 and 316-44 cells, APP signal is completely abolished. In 316-15 cells, although most amino acids of APP are retained, its localization to the Golgi apparatus is lost, and it diffusely exists in the cytoplasm. **c**, APP mutant cells become vulnerable to cellular toxicity induced by treatment with etoposide (0 or 20 μM, day 1) or LMNA siRNA (day 3). Graphs represent cell viability, with values indicating the data averaged to untreated Hela cells or Hela - control siRNA. Statistical analysis: ordinary one-way ANOVA with Bonferroni posthoc test for multiple comparisons. **d**, Western blot analysis of cleaved Gasdermin D and cleaved caspase-3. APP mutant cells exhibit an increase in cleaved Gasdermin D and cleaved caspase-3 following treatment with etoposide (0 or 20 μM, day 2) or LMNA siRNA (day 4). Uncropped immunoblots are provided in Supplemental Fig. 2b. **e**, Intracellular inflammatory response assessed for IL6 and IL1β upon treatment with etoposide (0 or 20 μM, day 2) or LMNA siRNA (day 4), using RT-qPCR. Values represent the data averaged to untreated Hela cells or Hela - control siRNA. APP mutant cells show an increased intracellular inflammatory response upon treatment with etoposide or LMNA siRNA. Statistical analysis: ordinary one-way ANOVA with Bonferroni posthoc test for multiple comparisons. **f**, Rescue effect of transfecting APP plasmid in APP mutant cells, assessed for IL1β, using RT-qPCR. APP WT transfection suppresses the inflammatory response in etoposide (20 μM, day 2) treated APP mutant cells, whereas this effect is absent in the mutant APP Swe/Ind transfection. Statistical analysis: ordinary one-way ANOVA with Bonferroni posthoc test for multiple comparisons. **g**, Reduction of endogenous APP exacerbates the etoposide-induced intracellular inflammatory response in iPSC-derived glutamatergic neurons (culture day 12, siRNA day 6). Graphs illustrate the intracellular inflammatory response evaluated for IL6 and IL1β upon etoposide treatment (0 or 20 μM, day 2), using RT-qPCR. Values represent the data averaged to control siRNA (untreated). Statistical analysis: ordinary one-way ANOVA with Bonferroni posthoc test for multiple comparisons. Data are mean ± s.e.m. The exact *P*-values of comparison are presented in the figures. \**P* < 0.05, \*\**P* < 0.01 and \*\*\**P* < 0.001.

To gain more insight to our results obtained from experiments using Hela cells, we next performed similar experiments using neurons differentiated from human iPSCs. Knockdown of endogenous APP resulted in a successful reduction of APP in these neurons (Fig. 2g). Following etoposide treatment, and in line with our previous results, the levels of inflammatory-related molecules, specifically IL6 and IL1β, were significantly increased in APP siRNA-treated neurons compared to control siRNA-treated neurons (Fig. 2g). In summary, we demonstrated that the loss of endogenous APP renders cells vulnerable to nuclear damage, as evidenced by experiments using Hela cells and human iPSCs-derived neuronal cells.

### APP disposes of nuclear waste resulting from nuclear damage

To further gain insight into the mechanism of APP’s protective role against inflammation and cell death caused by senescence-associated nuclear damage, we next analysed its subcellular localization in more detail. Co-expression of a plasmid expressing histone H2B and APP fused with a fluorescent protein at the N-terminus or C-terminus (respectively, TagRFP-H2B and APP-EGFP, and, EGFP-H2B and mRuby2-APP) resulted in the co-localization of nuclear-derived small dots of histone H2B and APP in the cytoplasm in both N-terminus and C-terminus fusion cases, following etoposide treatment (Fig. 3a). This indicated that full-length APP695, but not the cleaved form, primarily co-localizes with nuclear-derived small cytoplasmic dots of histone H2B upon senescence-associated nuclear damage. Similarly, endogenous APP also co-localized with nuclear-derived small dots of histone H2B following nuclear damage (Fig. 3b). Based on our RNA sequencing results (Extended data Fig. 1b,c) and accounting for the known functions of APP, we hypothesized that APP might dispose of nuclear waste generated by nuclear damage into the extracellular space. To challenge this hypothesis, we proceeded with the analysis of extracellular fluid using Hela cells and human iPSC-derived glutamatergic neurons. Nuclear damage led to an increase in nuclear-derived DNA components in the extracellular space; notably, analysis of the extracellular fluid from all APP mutant cell lines (316-15, 316-34, and 316-44) demonstrated significantly low amounts of these components (Fig. 3c). At the protein level, nuclear damage caused an increase in extracellular nuclear-derived histone H2B and Annexin A5 (ANXA5), which was markedly reduced in APP mutant cell lines (Fig. 3d). In particular, comprehensive proteomics analysis of the extracellular fluid revealed selective extrusion of nuclear components and intracellular retention of mitochondrial components in etoposide-treated WT Hela cells (Extended Data Fig. 2a,b). Conversely, in etoposide-treated APP mutant cell line (316-34), the extrusion of nuclear components into the extracellular fluid was significantly reduced in comparison to etoposide-treated WT cells (Fig. 3e). More, confocal microscopy analysis depicted abnormal structures adhering to the cell membrane in APP mutant cells (316-34) with senescence-associated nuclear damage, which were scant in WT Hela cells, and these abnormal structures contained nuclear-derived components (Fig. 3f, Extended Data Fig. 2c and Supplementary Movies 1,2). Similarly, knockdown of APP and treatment with etoposide in neurons differentiated from human iPSCs showed a high number of abnormal structures close to the plasma membrane (Extended Data Fig. 2d). Electron microscopy analysis revealed abundant extracellular disposal of nuclear-derived waste in WT Hela cells upon nuclear damage, whereas APP mutant cells (316-34) exhibited abnormal nuclear-derived waste adhering to the cell membrane (Fig. 3g). Intriguingly, live imaging revealed that APP mediates the extracellular disposal of nuclear waste leaked into the cytoplasm upon nuclear damage (Extended Data Fig. 2e and Supplementary Movies 3-6). In summary, we demonstrate that APP disposes of nuclear-derived waste out of the cell when the nucleus faces damaging stimuli.

**Fig. 3.**
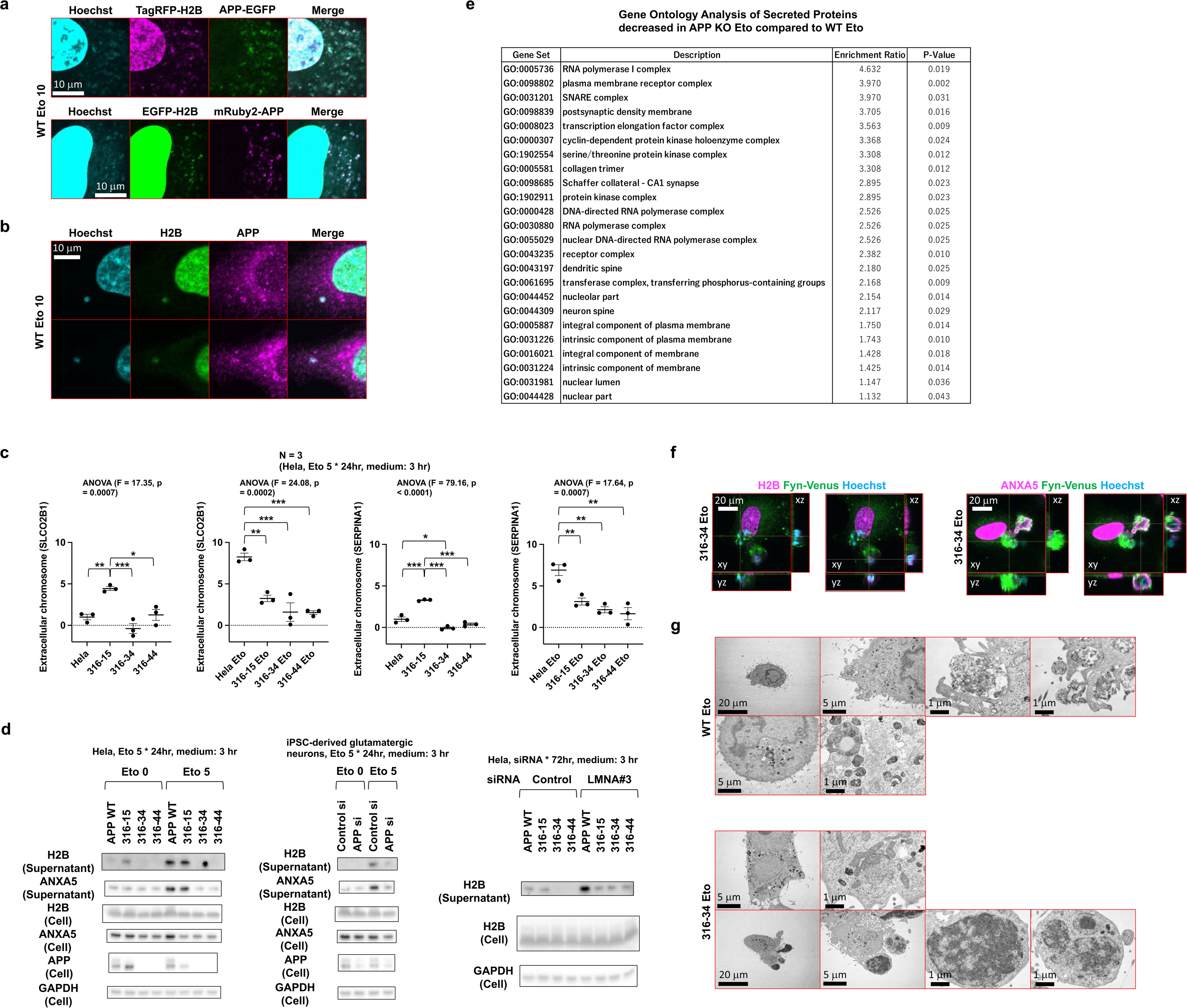
APP mediates extracellular disposal of nuclear waste. **a**, Co-localization of APP with nuclear-derived structures in the cytoplasm during nuclear damage (etoposide treatment 10 μM, day 1). Fluorescence images of APP-EGFP (C-terminal tag)/TagRFP-Histone H2B (N-terminal tag), or mRuby2-APP (N-terminal tag)/EGFP-Histone H2B (N-terminal tag) are presented. Co-localization with histone H2B was observed regardless of whether the fluorescent protein was bound to the C-or N-terminus of APP, suggesting that the full-length APP protein is primarily co-localized with histone H2B. **b**, Immunostaining of endogenous APP and Histone H2B. **c**, Quantification of extracellular DNA in cell culture supernatant using RT-qPCR. Cells were exposed to etoposide (0 or 5 μM) for 24 hrs, then the medium was washed, and the cells were cultured in serum-free medium for additional 3 hrs. The values from the Proteinase K-treated sample are subtracted from the values from the sample before the Proteinase K treatment. Statistical analysis: ordinary one-way ANOVA with Bonferroni posthoc test for multiple comparisons. **d**, Western blotting analysis of proteins in cell culture supernatant. Left, results from Hela cells and various APP mutant cell lines (316-15, 316-34, and 316-44) treated with etoposide. Middle, results from human glutamatergic neurons derived from iPSCs subjected to APP knockdown and etoposide treatment. WT Hela, APP mutant cells and human glutamatergic neurons were exposed to etoposide (0 or 5 μM) for 24 hrs, then the medium was washed, and the cells were cultured in serum-free medium for additional 3 hrs. Right, results from Hela cells and various APP mutant cell lines treated with LMNA siRNA (day 3). At day 3, the medium was washed, and the cells were cultured in serum-free medium for additional 3 hrs. Uncropped immunoblots are provided in Supplemental Fig. 3. **e**, DIA proteomics analysis of proteins in cell culture supernatant comparing Hela and APP mutant cells (316-34) upon etoposide treatment. Cells were exposed to etoposide (5 μM) for 24 hrs, then the medium was washed, and the cells were cultured in serum-free medium for additional 3 hrs. The table shows gene ontology analysis of decreased proteins (< 25%) in etoposide-treated APP mutant cells compared to etoposide-treated Hela cells. Nucleus-chromatin related proteins are reduced in the cell culture supernatant of APP mutant cells. **f**, Confocal fluorescence microscopy images of etoposide-treated (10 μM, 24 hrs) APP mutant cells (316-34), indicating the presence of abnormal structures adhering to etoposide-treated APP mutant cells (316-34). Membrane-targeted sequences of Fyn kinase were linked to the Venus fluorescent protein, allowing visualization of the plasma membrane in etoposide-treated APP mutant cells (316-34). **g**, Top, electron microscopy images of etoposide-treated (10 μM, 24 hrs) Hela cells, revealing the extracellular disposal of nuclear waste. Bottom, electron microscopy images of APP mutant cells (316-34) treated with etoposide (10 μM, 24 hrs), demonstrating the inability to dispose of nuclear waste, resulting in the accumulation of abnormal structures on the cell surface. Data are mean ± s.e.m. The exact *P*-values of comparison are presented in the figures. \**P* < 0.05, \*\**P* < 0.01 and \*\*\**P* < 0.001.

### Extracellular disposal of nuclear waste is impaired in mutant APP Swe/Ind

We further examined the relationship between the loss of protective effect against nuclear toxicity in mutant APP Swe/Ind and the extracellular disposal system of nuclear waste. As demonstrated in the previous section, APP WT successfully disposed of nuclear-derived waste into the extracellular space following etoposide treatment; however, this was not the case for APP Swe/Ind (Fig. 4a). Similarly, APP WT disposed of nuclear-derived waste into the extracellular space when cells were treated with LMNA siRNA, whereas this capability was impaired in APP Swe/Ind (Fig. 4b). Interestingly, in both etoposide and LMNA siRNA treatment, there was a significant increase in the extracellular release of C-terminal fragment of APP upon nuclear stress, suggesting that extracellular deposition of Aβ might be a consequence rather than a cause of cellular damage (Fig. 4a,b). We hypothesized that the difference in binding affinity to nuclear waste between APP WT and APP Swe/Ind may be the reason why APP WT can extracellularly excrete nuclear waste more efficiently than APP Swe/Ind. To test this, we performed immunoprecipitation experiments using FLAG-tagged APP. The results demonstrated that APP Swe/Ind’s capability of binding to nuclear - especially chromosome-derived proteins – was reduced when compared to APP WT (Fig. 4c). In summary, we unravel a novel role of APP, which is the disposal of nuclear-derived waste into the extracellular space following nucleus impairment; nevertheless, this intrinsic role is diminished in APP mutant Swe/Ind.

**Fig. 4.**
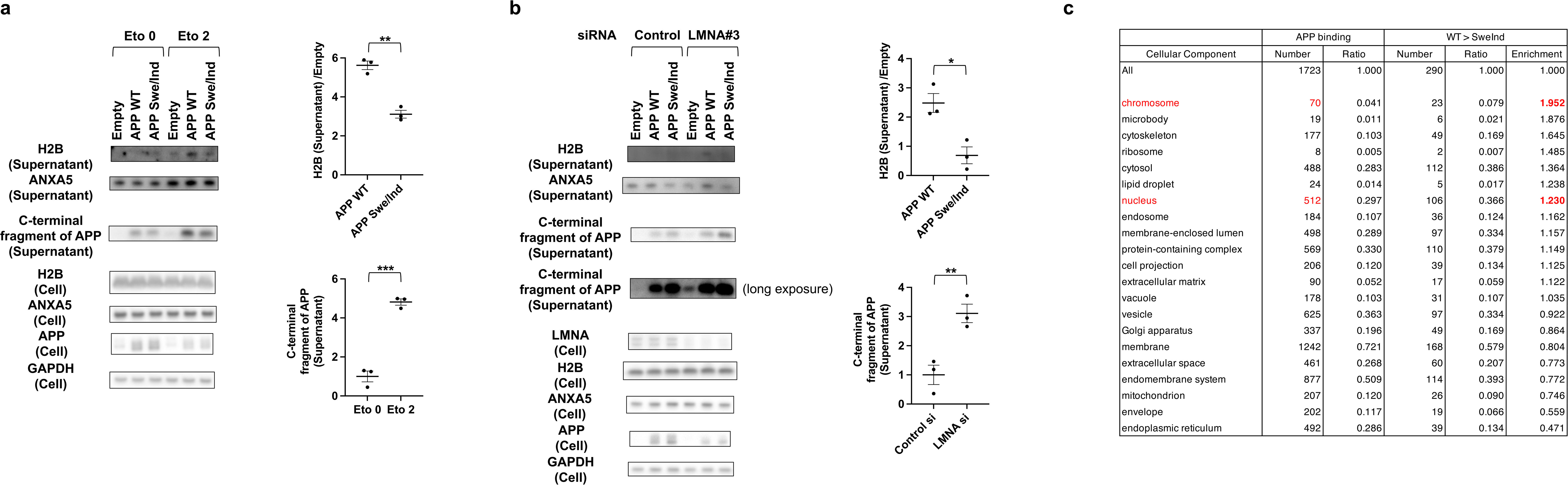
Nuclear waste disposal is impaired in APP Swe/Ind mutant. **a**, Western blot analysis was conducted on proteins present in cell culture supernatants. Results were obtained from HeLa cells overexpressing empty vectors, APP WT, and Swe/Ind mutant APP, respectively with or without etoposide treatment (0 or 2 μM, for 24 hrs). Uncropped immunoblots are provided in Supplemental Fig. 4a. Cells were exposed to etoposide, then the medium was washed, and the cells were cultured in serum-free medium for additional 3 hrs. The upper graph on the right side illustrates the relative amount of H2B protein, while the lower graph shows the relative amount of C-terminal fragment of APP in the supernatant. Statistical analysis: two-tailed, unpaired Student’s t-test. **b**, Western blot analysis was performed on proteins present in the cell culture supernatant. Results were obtained from HeLa cells overexpressing empty vectors, APP WT, and Swe/Ind mutant APP, respectively with or without LMNA siRNA treatment (day 3). Uncropped immunoblots are provided in Supplemental Fig. 4b. At day 3, the medium was washed, and the cells were cultured in serum-free medium for additional 3 hrs. The upper graph on the right side depicts the relative amount of H2B protein, while the lower graph illustrates the relative amount of C-terminal fragment of APP in the supernatant. Statistical analysis: two-tailed, unpaired Student’s t-test. \**P* < 0.05, \*\**P* < 0.01 and \*\*\**P* < 0.001. **c**, DIA-proteomics analysis of anti-FLAG immunoprecipitates from HeLa cells transfected with either empty vector, APP WT-FLAG, or Swe/Ind mutant APP-FLAG followed by etoposide treatment (5 μM, for 24 hrs). Enrichment of gene ontology terms in “cellular component” database is shown for APP-bound proteins (APP binding; APP WT / empty vector > 3) as well as for proteins which showed higher affinity to APP WT than to Swe/Ind mutant APP (WT>Swe/Ind; APP WT / APP SweInd > 2).

### Etoposide induces DNA damage, cell death and cytoplasmic histone accumulation in neurons of the mouse cortex

Having established the protective role of APP WT following nuclear insults using Hela cells and human iPSC-derived neurons, we proceeded with experiments in mice with the idea that APP WT, but not APP Swe/Ind, could also be protective in vivo. Because our hypothesis differs significantly from the existing Aβ hypothesis, it was difficult to use AD mouse models that were built on the Aβ hypothesis. High-dose etoposide was previously reported to induce DNA damage in various mouse strains, and increase the production of reactive oxygen species (ROS) in human glioblastoma cells leading to increased p53-mediated cellular apoptosis^20–22^. Thus, to investigate the protective role of APP, we utilized mice subjected to DNA damage following intraperitoneal administration of etoposide (Fig. 5a). Immunostaining of γH2AX foci revealed a significant increase in DNA damage in neurons of the cortex of the etoposide-treated mice in comparison to the control group (P < 0.0001, Fig. 5b,c). We further confirmed increased γH2AX in the etoposide-treated mice by performing western blotting of mouse brains (Fig. 5d). Next, to evaluate the impact of etoposide-induced DNA damage on cell viability within the mouse cortex, we conducted immunostaining of cleaved caspase-3, a key marker of apoptosis. We observed an increase in the levels of cleaved caspase-3 positive neurons in the etoposide-treated group compared to controls, indicative of enhanced cell death (Extended Data Fig. 3a,b).

**Fig. 5.**
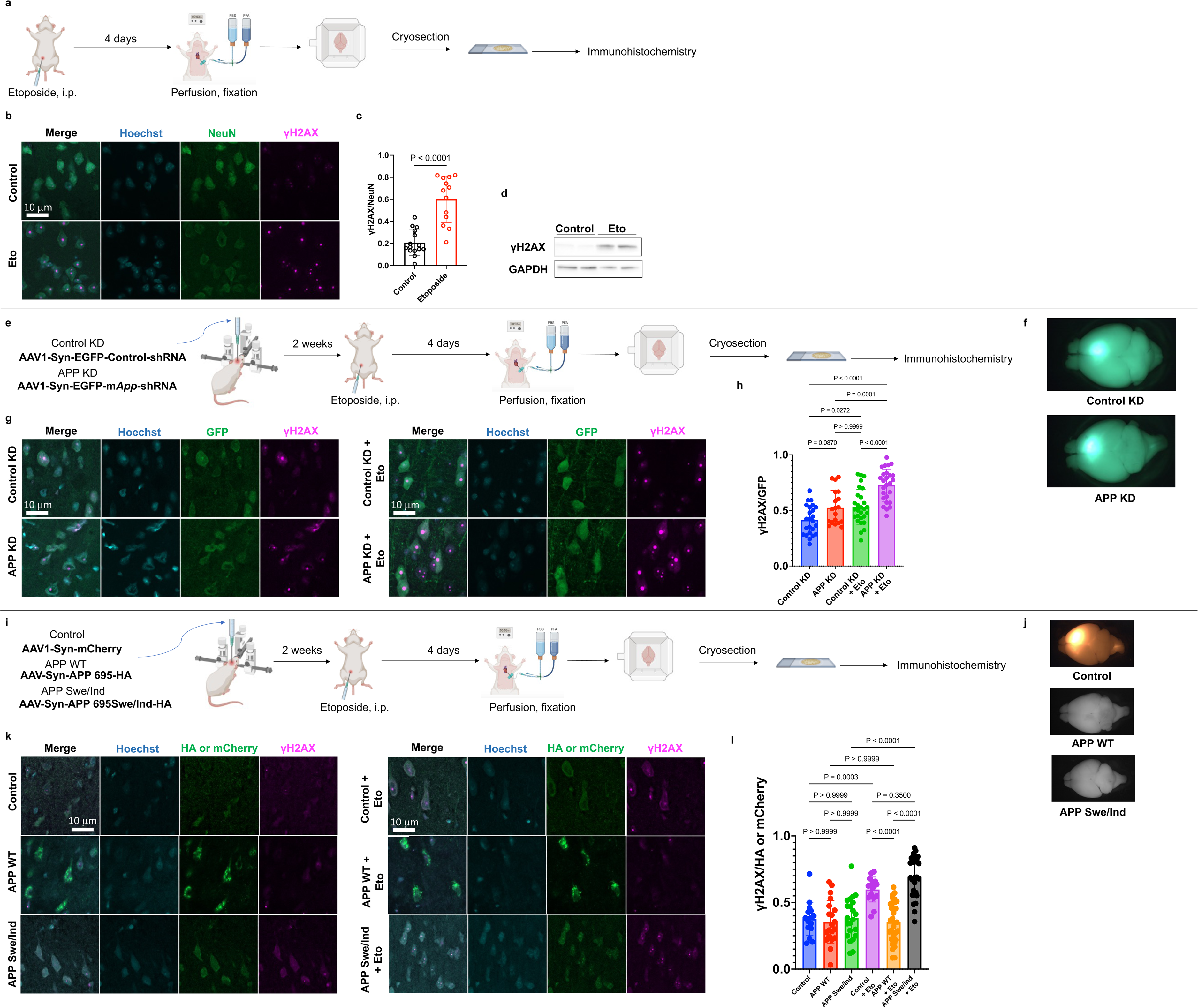
APP WT safeguards neurons challenged by DNA damage in vivo whereas APP Swe/Ind comes short. **a**, Scheme illustrating the experimental setup of DNA damage induction in the mouse cortex followed by perfusion fixation, cryosection, and immunohistochemistry; created with BioRender. i.p.: intraperitoneal injection. **b**, Representative immunofluorescence images of mouse cortex using Hoechst (nuclear marker, blue), NeuN (neuronal marker, green), and γH2AX (DNA DSBs marker, magenta) with and without (control) etoposide treatment. **c**, Quantification of γH2AX foci between control and etoposide-treated mice. Bars represent means; dots represent measurements for replicates (n = 6 mice per group, with 2-3 quantified images for each mouse). Statistical analysis: two-tailed, unpaired Student’s t-test (*P*-value is given in graph). **d**, Western blot analysis of γH2AX showing elevated levels in etoposide-treated mice relative to GAPDH, internal control. n = 2 mice per group. Uncropped immunoblot is provided in Supplemental Fig. 5b. **e**, Scheme illustrating the experimental setup of AAV injection of control KD and APP KD vectors in the mouse cortex followed by etoposide treatment, perfusion fixation, cryosection, and immunohistochemistry; created with BioRender. i.p.: intraperitoneal injection. **f**, Representative images showing successful AAV delivery with EGFP expression in control KD and APP KD mice. **g**, Representative immunofluorescence images of control KD and APP KD mice cortex using Hoechst (nuclear marker, blue), GFP (neuronal marker, green), and γH2AX (DNA DSBs marker, magenta) with and without (control) etoposide treatment. **h**, Quantification of γH2AX foci between control KD, APP KD, and both groups treated with etoposide. Bars represent means; dots represent measurements for replicates (n = 6 mice per group, with 2 quantified images for each mouse). Statistical analysis: ordinary one-way ANOVA with Bonferroni posthoc test for multiple comparisons (*P*-values for the posthoc tests are given in the graphs). **i**, Scheme illustrating the experimental setup of AAV injection of control, APP WT overexpression, and APP Swe/Ind overexpression vectors in the mouse cortex followed by etoposide treatment, perfusion fixation, cryosection, and immunohistochemistry; created with BioRender. i.p.: intraperitoneal injection. **j**, Representative images showing the brains delivered with control, APP WT and APP Swe/Ind overexpression AAV vectors. Note that APP WT and APP Swe/Ind showed no fluorescence because they were subcloned with the HA tag. **k**, Representative immunofluorescence images of control, APP WT and APP Swe/Ind mice cortex using Hoechst (nuclear marker, blue), HA or mCherry (neuronal marker, green) and γH2AX (DNA DSBs marker, magenta) with and without (control) etoposide treatment. **l**, Quantification of γH2AX foci between control, APP WT, and APP Swe/Ind mice and each group treated with etoposide. Bars represent means; dots represent measurements for replicates (n = 8 mice per group, with 2-3 quantified images for each mouse). Statistical analysis: ordinary one-way ANOVA with Bonferroni posthoc test for multiple comparisons (*P*-values for the posthoc tests are given in the graphs). Data are mean ± s.e.m.

It is well-known that UV irradiation or high dose toxicants can induce the release of nuclear histones into the cytoplasm following DNA-damage^23,24^. We then checked whether etoposide administration would induce profound damage to nuclear integrity, by investigating the presence of histone H2B in the cytoplasm as a hallmark of disrupted nuclear membrane integrity. Immunofluorescence staining confirmed the presence of cytosolic histone H2B in the etoposide-treated group, with a marked increase in mean and total cytosolic H2B (Extended Data Fig. 4a,b). This suggested that the nuclear contents leaked out of the nucleus in the etoposide group. Additionally, nuclear solidity, defined as the ratio of the measured area of the nucleus to the area of its convex hull shape (Extended Data Fig. 4a) was reduced in etoposide-treated compared to control mice (P < 0.0001, Extended Data Fig. 4c). In the etoposide-treated mouse cortex, increased DNA damage evidenced by elevated γH2AX foci formation, a significant increase in cell death indicated by high levels of cleaved caspase-3, and the presence of histone H2B in the cytoplasm of the mouse neurons highlight AD-like neurodegeneration and collapse of nuclear membrane integrity^25,26^. In sum, we established that etoposide treatment underscores a profound disruption of nuclear and cellular homeostasis within the mouse brain cortex.

### Knockdown of APP in the mouse cortical neurons exacerbates etoposide-induced neuronal damage

We next proceeded to investigate the intrinsic involvement of APP and its function in the mouse brain. Initially, we constructed two shRNA vectors, opting for shRNA#1 due to its superior knockdown efficiency of mouse APP in Neuro-2a cells, for subsequent in vivo experiments with mice (Extended Data Fig. 5a,b). We reduced APP expression in neurons through delivery of adeno-associated viral (AAV) vectors expressing shRNA against mouse *App* with GFP and subsequently performed immunostaining of the cortex sections (Fig. 5e). Two weeks post-injection, we confirmed successful AAV injection (Fig. 5f) and observed a clear reduction of APP in the GFP positive neurons of APP knockdown (KD) mice in comparison to the control KD group (Extended Data Fig. 5c,d).

Mice in each group were then treated with etoposide to induce nuclear damage, and the effects of APP KD after neuronal damage were evaluated. In the control KD group treated with etoposide, we noted a substantial increase in γH2AX foci compared to the control group without etoposide, consistent with the etoposide-induced DNA damage (Fig. 5g). In the APP KD group treated with etoposide, there was an increase in DNA damage, surpassing the levels observed in the control KD group treated with etoposide (F (3, 93) = 19.74, P < 0.0001, Fig. 5g,h). These findings indicate that the loss of physiological functions of APP has made neurons vulnerable to nuclear toxicity associated with DNA damage. Further, we compared DNA damage in the AAV injected and etoposide-treated APP KD group to the non-injected contralateral mouse cortex. Interestingly, DNA damage was higher in the AAV injected side (GFP positive neurons) in comparison to the GFP negative neurons (contralateral region, Extended data Fig. 5e,f). It should be mentioned that AAV injection itself increases DNA damage^27,28^. However, despite the increased DNA damage due to AAV, the DNA damage in APP KD group treated with etoposide was conspicuous when compared to the control KD group treated with etoposide.

We then sought to know about the consequences of APP KD on cell death in the context of etoposide-induced DNA damage. Immunostaining revealed a substantial increase in cleaved caspase-3 positive neuron levels in the etoposide-treated control KD group compared to the control KD group without etoposide (F (3, 60) = 11.10, P < 0.0001, Extended Data Fig. 6a,b). Interestingly, a higher proportion of cleaved caspase-3 positive neurons was observed in the etoposide-treated APP KD group, when compared to the etoposide-treated control KD group (Extended Data Fig. 6b). This is despite the fact that evidencing cell death in vivo remains a challenging assay, as apoptotic bodies are rapidly ingested by professional phagocytes such as microglia, monocytes and macrophages or more slowly by adjacent stromal cells such as fibroblasts and vascular smooth muscle cells^29^. This indicated that knockdown of APP caused increased apoptosis under genotoxic stress, as evidenced by higher cleaved caspase-3 positive neurons, and these results are in line with our cell experiments.

Consequently, we conducted immunofluorescence staining for histone H2B, and neurons with increased cytoplasmic histone H2B were observed in the etoposide-treated control KD group in comparison to the control KD group without etoposide. Furthermore, mice in the APP KD group treated with etoposide displayed a pronounced presence of cytoplasmic histone and abnormal nuclear solidity when compared to the control KD group treated with etoposide (F (3, 846) = 20.44, P < 0.0001, Extended Data Fig. 7a,b).

These results unequivocally demonstrate that APP KD substantially increases DNA damage, cell death and nuclear impairments compared to the control KD group under etoposide treatment. We demonstrate the consequences of APP loss of function in the context of nuclear damage and suggest that APP may hold substantial implications for conditions marked by neurodegeneration and DNA damage, such as AD.

### APP WT but not APP Swe/Ind alleviates the etoposide-induced DNA damage, cell death and cytoplasmic histone accumulation in vivo

Having evidenced that APP protects neurons from nuclear damage in mice, we next proceeded to test the hypothesis that APP WT but not APP Swe/Ind mutant would mitigate the etoposide-induced nuclear damage and cell death. To challenge this hypothesis, we overexpressed APP WT or APP Swe/Ind using AAV and intra-cortically injected AAV1-Syn-mCherry (control vector), AAV-Syn-APP 695-HA (APP WT overexpression) or AAV-Syn-APP 695Swe/Ind-HA (APP Swe/Ind overexpression) in a similar fashion as we did for APP KD investigations (Fig. 5i). We confirmed APP overexpression two weeks post-injection (Fig. 5j) and measured APP fluorescence in the control, APP WT and APP Swe/Ind groups (Extended Data Fig. 8a,b). Compared to the control group, both APP WT and APP Swe/Ind overexpression were confirmed by an increase in APP fluorescence (F (2, 316) = 90.72, P < 0.0001, Extended Data Fig. 8b).

Next, mice were subjected to etoposide treatment to induce nuclear damage. In the control group treated with etoposide, as expected, we observed a significant increase in the number of γH2AX foci, indicative of DNA damage (Fig. 5k). However, in the APP WT overexpression group treated with etoposide, the number of γH2AX foci was noticeably reduced compared to the control group treated with etoposide (Fig. 5l). Additionally, we compared DNA damage in the AAV injected region to the contralateral mouse cortex, and results confirmed the reduction of γH2AX foci in APP WT overexpression (Extended data Fig. 8c,d). Strikingly, in the APP Swe/Ind overexpression group treated with etoposide, there was no discernible reduction in γH2AX foci, and the damage level resembled that of the control group treated with etoposide (Fig. 5l, Extended data Fig. 8c,d). Further, quantitative analysis validated these observations as the etoposide-treated APP WT group exhibited a statistically significant reduction in the number of γH2AX foci compared to both the etoposide-treated control and APP Swe/Ind groups (F (5, 127) = 24.68, P < 0.0001, Fig. 5l). This confirmed again the crucial role of APP in mitigating DNA damage and nuclear impairment, and this protective effect was not observed with the APP Swe/Ind mutation.

Next, to assess the impact of APP WT and APP Swe/Ind overexpression on cell death in the presence of etoposide-induced DNA damage, we utilized cleaved caspase-3 as marker for apoptosis. In the control group subjected to etoposide treatment, there was an increase in cleaved caspase-3 levels compared to the control group. Notably, in the APP WT group treated with etoposide, the levels of cleaved caspase-3 were lower, albeit not significantly, than in the control group treated with etoposide (Extended Data Fig. 9a,b). However, in the etoposide-treated APP Swe/Ind overexpression group, cleaved caspase-3 levels were significantly increased in comparison to the etoposide-treated control group (Extended Data Fig. 9a,b). This indicated the role of APP WT, but not mutant APP, in reducing cell death upon DNA damage.

Next, we asked about nuclear solidity and cytosolic histones following etoposide administration and APP WT or APP Swe/Ind overexpression. In the control group treated with etoposide, we observed a notable increase in cytoplasmic histone H2B, indicating disrupted nuclear membrane integrity (Extended Data Fig. 10a,b). Interestingly, in the APP WT group treated with etoposide, the extent of cytosolic histone H2B was significantly reduced compared to the etoposide-treated control group (Extended Data Fig. 10a,b). By contrast, the APP Swe/Ind group treated with etoposide failed to reduce cytosolic H2B, and quantitative analysis of nuclei shape further demonstrated abnormal nuclear solidity in the etoposide-treated APP Swe/Ind group in comparison to the etoposide-treated APP WT group, indicative of nuclear anomality and fragility (F (5, 968) = 18.02, P < 0.0001, Extended Data Fig. 10b). It is interesting that APP WT confers protection against DNA damage, cell death, and disruption of nuclear membrane integrity in both cellular and mice experiments. Contrarywise, APP mutant Swe/Ind does not provide such protective functions at the cellular level or in vivo.

### Reduction of APP and accumulation of DNA damage is marked in human AD neurons

Lastly, we conducted investigations using postmortem human AD brains. We focused on the anterior cingulate cortex, which is part of the cerebral cortex and is frequently disturbed in AD. Results indicated a higher deposition of histones in the cytoplasm of AD brain neurons compared to control brains (Fig. 6a). Additionally, through analysing numerous cell nuclei, we revealed abnormal nuclear morphology in AD neurons, mirroring our findings in etoposide-treated mice (Fig. 6a). Further, AD neurons exhibited a significant increase in γH2AX foci, compared to control and Parkinson’s disease (PD) brains, indicative of increased DNA damage (Fig. 6b). Moreover, western blot analysis demonstrated an elevation of γH2AX and cleaved PARP1 in AD neurons (Fig. 6c). Interestingly, it is worth mentioning that APP levels were reduced in AD brains, suggesting that the same cascade that occurred in cells and mice with APP knocked down could occur in human AD brains (Fig. 6c). To investigate whether the reduction of APP was due to a decrease in intraneuronal APP or simply a result of neuronal loss in the brain, we then performed immunostaining for APP using control and AD brains. As a result, we observed a significant decrease in the levels of APP in AD neurons compared to control and PD neurons (Fig. 6d). In summary, a decrease in APP and an increase of DNA damage were observed in human neurons affected by AD. These findings suggest a potential implication of the interplay between reduced APP and elevated DNA damage in the neurodegenerative mechanisms observed in AD. These findings highlight a pivotal role of APP in safeguarding cellular homeostasis in the presence of genotoxic stress and emphasize the functional significance of APP mutations in compromising its protective capabilities. Our study advances the understanding of APP’s role in cellular defense mechanisms and highlights its potential implications in AD and other pathological conditions marked by nuclear damage and cellular disruption (Extended data Fig. 11).

**Fig. 6.**
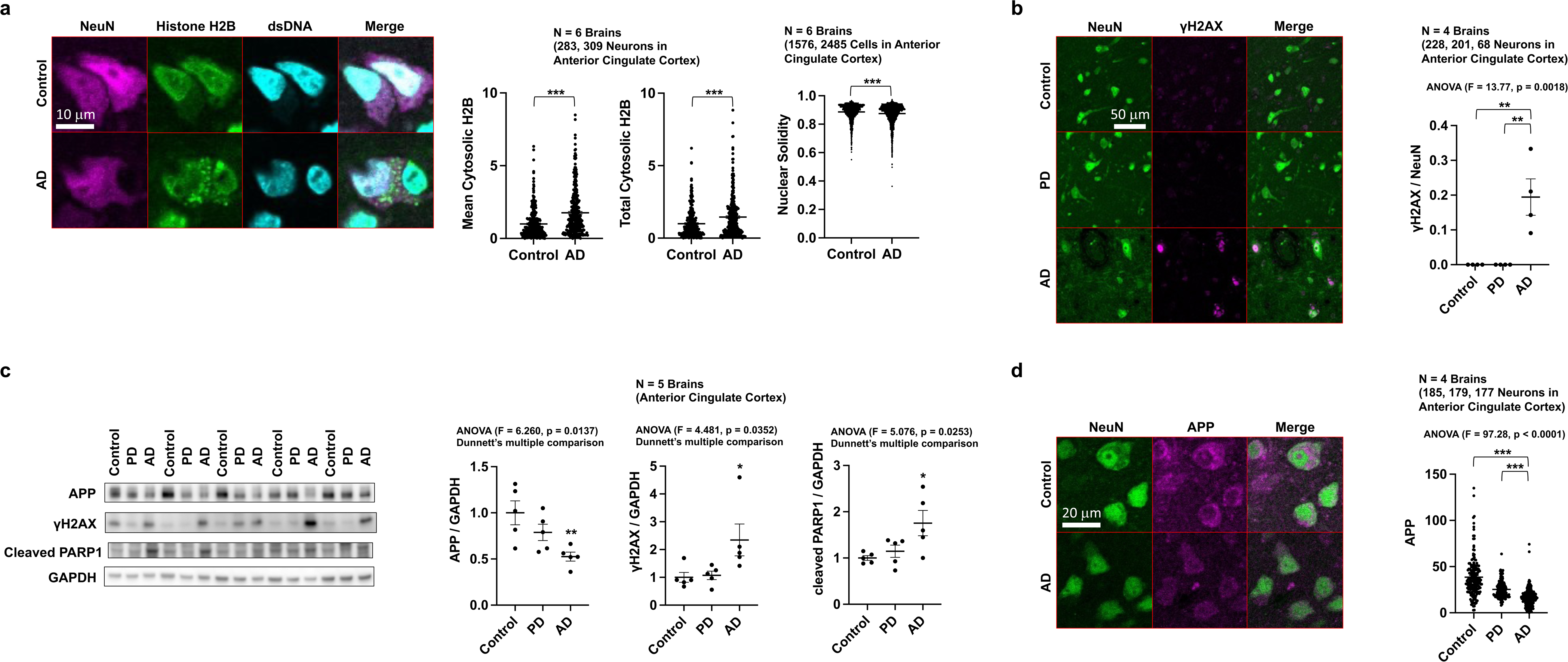
Increased nuclear waste and decreased APP in human AD neurons. **a**, Left, confocal images of cytoplasmic accumulation of nuclear-derived waste in neurons of human AD brains. Right, quantification of Histone H2B signals in the cytoplasm and nuclear solidity in neurons. Abnormal nuclear morphology is evidenced in neurons of human AD brains. Statistical analysis: two-tailed, unpaired Student’s t-test. **b**, Left, confocal images of human AD brains stained using anti-γH2AX antibody, a marker of nuclear damage, show increased DNA damage in neurons of human AD brains. Right, quantification of γH2AX foci. Statistical analysis: ordinary one-way ANOVA with Bonferroni posthoc test for multiple comparisons. **c**, Western blotting analysis of human AD brain lysates. Human AD brains exhibit elevated DNA damage markers and decreased APP levels. Statistical analysis: Dunnett’s multiple comparison. Uncropped immunoblot is provided in Supplemental Fig. 6. **d**, Left, immunohistochemistry of human AD brains using anti-APP antibody shows reduced levels of APP per neuron in human AD brains. Right, quantification of APP levels in neurons. Statistical analysis: ordinary one-way ANOVA with Bonferroni posthoc test for multiple comparisons. Data are mean ± s.e.m. The exact *P*-values of comparison are presented in the figures, respectively. \**P* < 0.05, \*\**P* < 0.01 and \*\*\**P* < 0.001.

## Discussion

The accumulation of Aβ in the brains of individuals with AD, the identification of APP as one of the causative genes in familial AD, and the evidence of Aβ cytotoxicity, may have contributed to the prevailing belief that Aβ accumulation is the primary cause of AD^30^. However, after more than 30 years of intensive research, AD remains a challenging and therapeutically elusive condition presenting significant medical and societal concerns, with a growing number of patients affected due to the aging population. The Aβ theory of AD has been beset by numerous investigations, huge cost investments, and limited clinical treatments^2,3,31^, and understanding of the disease’s broader picture will ultimately lead to treatments. For years, researchers have focused on AD’s telltale amyloid plaques – complex tangles of Aβ, cleaved product of APP frequently found in the brains of patients with neurodegenerative disorders, but controversially with an exceedingly high prevalence in the brains of neurotypicals^32,33^. In exploring ectopic mitochondrial DNA in the cytosol and its implications for PD neurodegeneration^34^, we developed a hypothesis regarding a possible protective role of APP in DNA damage. Recent findings suggest that APP might be involved in safeguarding genomic integrity, and functions as an important component for the development of the nervous system relating to synapse structure and function, as well as in neuronal migration or adhesion^11,35,36^. However, to date, the role of APP in AD development and progression remains unclear, and it is not well-understood whether it contributes to these aspects of the disease.

Here, we unravel a new role of APP in the context of neurodegenerative disorders like AD. The exploration of APP’s potential to rescue DNA damage, inflammation, and cell death in both in vitro and in vivo models holds significant relevance, as it provides compelling evidence for its neuroprotective role in combating AD-associated pathophysiology. Indeed, APP WT exhibited various protective effects, including the inhibition of inflammatory responses and cell death, following nuclear damage induced by etoposide treatment or LMNA KD. Additionally, in cell experiments, APP WT facilitated the disposal of nuclear-derived waste materials outside the cell, thus maintaining nuclear and cellular homeostasis. Intriguingly, these protective effects were significantly diminished in an AD-associated mutant form of APP, APP Swe/Ind. Our findings were further supported by in vivo experiments in mice, where reducing endogenous mouse APP exacerbated neuronal damage induced by intraperitoneal administration of etoposide. Conversely, overexpression of human APP WT in mice reduced neuronal damage caused by etoposide, whereas this neuroprotective effect was not noticeable in APP Swe/Ind. We hypothesized that the absence of protective activity in APP Swe/Ind may be attributed to a diminished binding capability to nuclear waste, and this was confirmed through immunoprecipitation analysis. Further, in postmortem human AD brains, a significant increase in nuclear damage in neurons was observed, accompanied by abnormal nuclear morphology. More, the levels of APP per neuron were significantly reduced in AD compared to neurotypical control brains, and we speculate that low levels of APP in AD patients could lead to APP incapacity to protect the cells and organism from inevitable neurodegeneration.

The cumulative effects of DNA damage, inflammation, and subsequent cell death in AD pathogenesis have gained particular attention in recent research. Growing evidence suggests a substantial interplay between these factors and the progression of AD pathology. Recently, Mathys et al. generated a single-cell transcriptomic atlas of the aged human prefrontal cortex covering 2.3 million cells from postmortem human brain samples of 427 individuals with varying degrees of AD pathology and cognitive impairment^37^. Interestingly, they identified AD-associated alterations shared between excitatory neuron subtypes and revealed a coordinated increase of the cohesin complex and DNA damage response factors in excitatory neurons and oligodendrocytes. Further, Dileep et al. showed that following DNA insults in a neurodegeneration-model mouse, the 3D genome is altered with gene fusions particularly enriched in excitatory neurons with DNA damage repair and senescence gene signatures showing transcriptional changes in synaptic, neuronal development, and histone genes^5^. Furthermore, neurons burdened with DSBs following etoposide treatment exhibited disruption of RAD21 and Lamin B1, two structural proteins involved in DSB repair and 3D genome organization, in addition to increased senescence-like and immune gene expression signatures. In our cell culture experiments, C-terminal fragment of APP was extracellularly excreted following nuclear damage, suggesting that extracellular accumulation of Aβ may not be the cause of cell damage, but rather, could be a consequence of neurons and nuclei burdened by DNA damage. Nevertheless, it might be premature to completely abandon the Aβ hypothesis, as it is possible that both mechanisms occur in such a way that nuclear damage leads to extracellular accumulation of Aβ, which in turn exacerbates cell toxicity, leading to a noxious cycle. Additionally, it is safe to speculate that an altered processing of APP, and in turn a modification of its functions, could negatively affect the response of neurons to DNA damage in AD brain, thus leading to chronic accumulation of DNA damage that could significantly contribute to the neuronal loss characteristics of AD pathology^38^. Several previous reports are consistent with our findings, suggesting that persistent DNA DSBs in neurons are an early pathological hallmark of AD, and that DSBs may be upstream to the cascade of events leading to toxicity in neurodegenerative diseases^25,26,39–42^. Nuclear damage is closely associated with aging, and our findings also suggest the presence of abnormal aging processes underlying AD. Yujun et al. showed impaired neuronal bioenergetics and neuroinflammation in the progression of AD, and nicotinamide riboside treatment reduced pyrin domain containing 3 (NLRP3) inflammasome expression, DNA damage, apoptosis, and cellular senescence in the AD mouse brains^43^. Furthermore, protective effects against nuclear damage have been reported for microtubule associated protein tau (MAPT), another crucial protein in AD known for its accumulation^44^. Alternatively, the potential of senescent cell clearance-senolysis-to ameliorate phenotypes associated with aging and age-related diseases has been documented^45^, and AD could also be improved through senolysis^46^.

One limitation of our study is that behavioral assessment as originally intended, particularly comparing the potential of APP WT to rescue altered behaviors associated with DNA damage-thus neurodegeneration and senescence^5,25,37,47^-in contrast to APP Swe/Ind, could not be conducted due to the high toxicity of etoposide on mice. The use of a high etoposide dose was pivotal to simulate conditions reflecting nuclear insults, and mirroring aspects of AD pathology. In the future, integrating complementary methodologies or models that offer avenues to conduct behavioral analyses could enhance our understanding of the interplay between APP variants and AD-associated behavioral changes.

Overall, while our report is groundbreaking, translating these findings into therapeutic strategies poses significant challenges. It is crucial to first examine the relationship between our hypothesis and extracellular Aβ deposits. The pursuit of interventions to suppress aging is also an active area of research, although it remains a complex task. Consequently, a promising therapeutic development strategy might involve enhancing pathways implicated in efficient degradation and disposal of nuclear waste materials, and we believe continued research efforts will help advance our understanding of AD and ultimately pave the way for innovative treatments.

## Methods

### Ethics Statement

All animal experiments were performed in compliance with the protocol reviewed by the Institutional Animal Care and Use Committee and approved by the President of Niigata University (#SA00656). Informed consent was obtained from the participants of human studies. The study using postmortem brain samples from human subjects was approved by the Ethical Review Boards of Niigata University (#2019-0272). Patient profiles are described in Supplementary Table 1.

### Cell Lines

HeLa cells (RCB0007) were derived from cervical cancer cells from a female and were provided by the RIKEN BRC through the National Bio-Resource Project of the MEXT/AMED, Japan. HeLa cells were cultured in DMEM (Nacalai Tesque, Kyoto, Japan) supplemented with 10% FBS (BioWest, Nuaillé, France) and 1% penicillin-streptomycin solution (Wako, Osaka, Japan) at 37°C under 5% CO_2_. For human iPSC-derived glutamatergic neurons, iCell GlutaNeurons (Fujifilm, Osaka, Japan) were used and grown in iCell Neural Base Medium supplemented with iCell Neural Supplement B and iCell Nervous System Supplement (Fujifilm) at 37°C in 5% CO_2_ atmosphere. APP-deficient or mutant HeLa cells were generated using the CRISPR-Cas9 system. The APP gRNA sequence was cloned into pSpCas9 (BB)-2A-Puro (PX495) v.2.0 (plasmid #62988; Addgene, Cambridge, MA, USA) using the method described by Ran et al.^48^. The overexpression of the plasmid DNA in HeLa cells was performed using Lipofectamine 3000 transfection reagent (Thermo Fisher Scientific, Waltham, MA, USA) according to the manufacturer’s instructions. After treatment with 2 μg/ml of puromycin for 48 hrs, cells were grown and passaged in medium without puromycin for 7 days and then spread in four 96-well plates at 0.75 cell/100 μl. Single colonies were screened by western blotting and genome sequencing, and reliable mutants were selected for establishment. The depletion or mutation of APP in the stable cell line was again verified and monitored using western blotting and genome sequencing.

### siRNA and Plasmid Transfection

The expression of genes of interest was silenced in iCell GlutaNeurons using Accell siRNAs in Accell siRNA delivery media (Dharmacon, Lafayette, CO, USA) according to the manufacturer’s instructions. Accell siRNAs were used at a final concentration of 1 μM unless otherwise stated. Cells were subjected to the subsequent experiments 5∼6 days after siRNA transfection in iCell GlutaNeurons, unless otherwise stated. The expression of genes of interest was silenced in HeLa cells using Stealth RNAi siRNA (Thermo Fisher Scientific) in Lipofectamine RNAiMAX Transfection Reagent (Thermo Fisher Scientific) according to the manufacturer’s instructions. Cells were subjected to the subsequent experiments 3∼5 days after siRNA transfection in HeLa cells, unless otherwise stated. The siRNAs used in the present study are listed in the Supplementary Table 2. The overexpression of the plasmid DNA in HeLa cells was performed using Lipofectamine 3000 transfection reagent (Thermo Fisher Scientific) according to the manufacturer’s instructions.

### Western Blotting

Total protein extracts were prepared in lysis solution (50 mM Tris, pH 8.0, 150 mM NaCl, 1% (v/v) NP40, 0.5% (w/v) deoxycholate, and 0.1% (w/v) SDS). Protein concentrations were determined using the BCA protein assay kit (Thermo Fisher Scientific). Protein extracts were separated in SDS-PAGE gels and transferred to a PVDF membrane (GE Healthcare, Chicago, IL, USA). Western blotting was performed according to standard protocols. For western blotting of the cell culture supernatant, cells were washed with PBS thrice and placed in serum-and phenol red–free DMEM, and conditioned media was collected after 3 hrs. Conditioned medium was centrifuged at 1,000g at 4°C for 5 min to remove cell debris and was used for further analysis. The details of the antibodies used in the present study are listed in the Supplementary Table 3.

### Cell Survival Assay

Cell viability was measured using a WST-8 assay (Cell Counting Kit-8, Dojindo, Kumamoto, Japan).

### RNA Isolation and RT-qPCR

Total cellular RNA was isolated using TRIzol reagent (Thermo Fisher Scientific) and Direct-zol RNA kit (Zymo Research, Irvine, CA, USA) according to the manufacturer’s instructions. The cDNA templates were synthesized from the purified RNA using the PrimeScript 1st strand cDNA Synthesis Kit (Takara Bio, Kusatsu, Japan) with oligo (dT)_20_ primers. RT-qPCR was performed using TB Green Premix Ex Taq II (Takara Bio) and analysed in a Thermal Cycler Dice Real Time System Lite (Takara Bio). The PCR primers used in the present study are listed in the Supplementary Table 4.

### Immunofluorescence Labelling of cells and human brain sections

Cells were washed twice with PBS and fixed with 4% paraformaldehyde (PFA) in PBS for 10 min at RT. The cells were washed three times with PBST for 5 min each, then permeabilized with PBS containing 0.2% (w/v) Triton X-100 for 10 min. The cells were washed three times with PBST for 5 min each and incubated with 2% (w/v) BSA in PBST for 30 min. The cells were incubated with primary antibodies in 2% (w/v) BSA in PBST for 1 hr at RT. The cells were washed three times with PBST for 5 min each and incubated with secondary antibodies diluted in 2% (w/v) BSA in PBST for 1 hr at RT. The cells were washed three times for 5 min each with PBST. When staining the nuclei with Hoechst, the cells were incubated with Hoechst 33258 (Dojindo) at the same time as the secondary antibodies.

For the labelling of endogenous histone, cells were washed twice with PBS and fixed with ice-cold methanol for 10 min at 4°C. The cells were washed three times with PBST for 5 min each, then permeabilized with PBS containing 0.2% (w/v) Triton X-100 for 10 min. The cells were washed three times with PBST for 5 min each and incubated with 2% (w/v) BSA in PBST for 30 min. The cells were incubated with a primary antibody against H2B diluted in 2% (w/v) BSA in PBST for 1 hr at RT. The cells were washed three times with PBST for 5 min each and incubated with a secondary antibody diluted in 2% (w/v) BSA in PBST for 1 hr at RT. The cells were then washed three times for 5 min each with PBST. If other molecules were to be stained, the staining was again performed from here in the same manner as for the primary and secondary antibodies.

For the imaging using postmortem brain tissues from human subjects, dewaxed paraffin sections (5 μm thickness) of the anterior cingulate cortex were incubated in 10 mM sodium citrate buffer (pH 6.0) at 121°C for 15 min in an autoclave (HA-300MIV, HIRAYAMA, Kasukabe, Japan). After the samples were allowed to cool gradually to RT, the sections were washed three times with PBS for 5 min each and incubated with 2% (w/v) BSA in PBS for 30 min. The sections were incubated with primary antibodies in 2% (w/v) BSA in PBS for 1 hr at RT. The sections were washed three times with PBS for 5 min each and incubated with secondary antibodies diluted in 2% (w/v) BSA in PBS for 1 hr at RT. The sections were washed three times with PBS for 5 min each and mounted with the SlowFade Gold antifade reagent (Thermo Fisher Scientific).

For the imaging of endogenous histone using postmortem brain tissues from human subjects, dewaxed paraffin sections (5 μm thickness) of the anterior cingulate cortex cut in the axial orientation were incubated in 10 mM sodium citrate buffer (pH 6.0) at 121°C for 15 min in an autoclave (HA-300MIV, HIRAYAMA). After the samples were allowed to cool gradually to RT, the sections were then immersed in methanol for O/N at 4°C. The sections were washed three times with PBS for 5 min each and incubated with 2% (w/v) BSA in PBS for 30 min. The sections were incubated with primary antibodies in 2% (w/v) BSA in PBS for 1 hr at RT. The sections were washed three times with PBS for 5 min each and incubated with secondary antibodies diluted in 2% (w/v) BSA in PBS for 1 hr at RT. If other molecules were to be stained, the staining was performed in the same manner as for the primary and secondary antibodies. Finally, the sections were washed three times with PBS for 5 min each and mounted with the SlowFade Gold antifade reagent (Thermo Fisher Scientific).

Immunofluorescence images were obtained using an A1R+ confocal microscope (Nikon Solutions, Tokyo, Japan). The details of the antibodies used in the present study are listed in the Supplementary Table 3.

### Immunoprecipitation and DIA proteome analysis

For immunoprecipitation of FLAG-tagged APP, HeLa cells cultured in a 10 cm dish were transfected with the plasmid expressing either empty vector, APP WT or APP Swe/Ind, tagged with 3x FLAG. One day after transfection, the medium was changed to DMEM containing 5 µM etoposide, and cells were cultured for another 24 hrs. Di (N-succinimidyl) 3,3’-dithiodipropionate (DSP, Tokyo Chemical Industry, Tokyo, Japan)-crosslinking immunoprecipitation (IP) was performed according to a previously described method with some modifications^49^. In brief, cells were rinsed twice with pre-warmed PBS and cross-linked with 0.1 mM DSP in PBS at 37°C for 30 min in a CO_2_ incubator. Cells were rinsed once with PBS, incubated in STOP solution (20 mM Tris-HCl pH 7.4 in PBS) at RT for 15 min and rinsed twice with ice-cold PBS. Next, 750 μL of ice-cold IP buffer (25 mM Tris-HCl pH 7.5, 300 mM NaCl, 1 mM EDTA, 0.5% NP-40, 1% TritonX-100, 5% glycerol with cOmplete Mini EDTA-free (Merck, Darmstadt, Germany)) was added and the cells were scraped off with a scraper, transferred to a 1.5 mL microtube and incubated on ice for 10 min to complete cell lysis. The lysates were centrifuged at 20,000g for 10 min at 4°C. Protein concentration of the supernatant was measured using BCA protein assay (Thermo Fisher Scientific) and adjusted to 2 mg/ml with IP buffer. Total 2 mg lysate (1 ml) was incubated with 10 µl of anti-FLAG M2 Magnetic Beads (Merck), which was pre-washed twice with IP buffer, at 4°C on a rotating wheel for 2 hr. DynaMag-2 Magnet (Thermo Fisher Scientific) was used for collecting the beads. The beads were then washed three times with 1 ml of IP buffer followed by once with 50 mM Tris-HCl (pH 8.0). Samples were digested on beads with trypsin (500 ng of Trypsin Platinum, Promega, Madison, WI, USA) in 100 µl of 50 mM Tris-HCl (pH 8.0) containing 2% lauryl maltose neopentyl glycol and 7.5 mM CaCl_2_ at 37°C for 15 hrs. Digested products were then reduced and alkylated with 10 mM tris (2-carboxyethyl) phosphine hydrochloride (TCEP) and 40 mM chloroacetamide, respectively. After acidification with final 5% trifluoroacetc acid (TFA), samples were desalted using reversed phase spin column (GL-Tip SDB, GL Science, Tokyo, Japan) followed by centrifugal evaporation and resuspension in 0.1% TFA-0.01% decyl maltose neopentyl glycol (DMNG). For LC-MS/MS analysis, the peptides were fractionated on 75 μm × 12 cm nanoLC column packed with C18 particle (3 µm; Nikkyo Technos Co., Ltd., Tokyo, Japan) by eluting with a linear gradient of 6% to 75% solvent B over 70 min at a flow rate of 200 nl/min (solvent A = 0.1% formic acid; solvent B = 0.1% formic acid in 80% acetonitrile) with the use of UltiMate 3000 RSLCnano LC System (Thermo Fisher Scientific). The Q Exactive HF-X (Thermo Fisher Scientific) mass spectrometer was operated in data-independent acquisition mode. MS1 spectra were collected in the range of 495–745 m/z at a resolution of 30,000 using an Automated Gain Control (AGC) target value of 3 × 10^6^ ions with a maximum injection time for 55 msec. MS2 spectra were collected in the range of > 200 m/z at a resolution of 30,000 using an AGC target value of 3 × 10^6^ ions and the automatic maximum ion injection times. Sixty data-independent acquisition (DIA) windows with isolation width of 4 Da were ranged from 500 to 740 m/z. The normalized collision energy was set to 23%. All DIA raw data was processed with DIA-NN (ver. 1.8)^50^ using library-free search mode against human UniProt sequences (https://www.uniprot.org/; accessed 7 March 2023). Enzyme specificity was set to “trypsin”. Precursor m/z range was set from m/z 490 to 75. Carbamidomethylation of cysteine was set as a fixed modification. Enrichment analysis of mass spectrometry data was performed using WEB-based Gene SeT AnaLysis Toolkit (https://www.webgestalt.org) with the setting of Over-Representation Analysis using “cellular component” of gene ontology as the database.

### Electron Microscopy

Samples were fixed with 2% PFA and 2% glutaraldehyde in 0.1 M phosphate buffer, pH 7.4, at 37°C then placed in a refrigerator for 30 min. The samples were fixed in 2% glutaraldehyde in 0.1 M phosphate buffer overnight at 4°C. The samples were washed three times with 0.1 M phosphate buffer for 30 min and postfixed with 2% osmium tetroxide in 0.1 M phosphate buffer at 4°C for 1 hr. The samples were dehydrated in graded ethanol solutions, transferred to resin (Quetol-812, Nisshin EM, Tokyo, Japan) and polymerized at 60°C for 48 hrs. The polymerized resins were cut into ultrathin 70-nm sections with a diamond knife using an ultramicrotome (Ultracut UCT, Leica Microsystems, Wetzlar, Germany) and mounted on copper grids. The sections were stained with 2% uranyl acetate at RT for 15 min and washed with distilled water, followed by secondary staining with a lead staining solution (Merck) at RT for 3 min. The grids were observed under a transmission electron microscope (JEM-1400Plus, JEOL, Tokyo, Japan) at an acceleration voltage of 100 kV, and images were recorded using a CCD camera (EM-14830RUBY2, JEOL).

### RNA Sequencing

RNA sequencing was performed by Macrogen Japan (Kyoto, Japan). Libraries were prepared using the TruSeq RNA Sample Prep Kit v2 (Illumina, San Diego, CA, USA) according to the manufacturer’s guidelines. The libraries were sequenced on a NovaSeq 6000 instrument with 100-bp paired-end reads (Illumina).

### Comet assay

DNA damage in cellular nucleus was measured using an OxiSelect Comet Assay Kit (Cell Biolabs, San Diego, CA, USA).

### DIA proteomics analysis of proteins in cell culture supernatant

To collect proteins secreted into serum-free medium, cells were seeded on 10 cm dish, then treated with or without 5 µM etoposide for 24 hrs. Subsequently, cells were washed with PBS thrice and placed in serum-and phenol red–free DMEM, and conditioned media was collected after 3 hrs. Conditioned medium was centrifuged at 1,000g at 4°C for 5 min to remove cell debris and filtered through 22 µm filter. Samples were concentrated using Vivaspan 20 with a 3000 MWCO (Sartorius Stedim Biotech GmbH, Goettingen, Germany) as per the manufacturer’s instructions. Protein samples were subjected to reduction and alkylation with 2.5 mM TCEP (Thermo Fisher Scientific) and 12.5 mM iodoacetamide (IAA) (Merck), respectively. After acetone precipitation of proteins, the pellets were sonicated in 100 mM HEPES pH8.0, followed by digestion with trypsin at a protein ratio of 1:50 at 37°C for 16 hrs. Resulting protein digests equivalent to 5 µg were desalted using StageTip (SDB-XC, 3M, St. Paul, MN, USA)^51^. The desalted peptides were injected (200 ng/sample) into a pre-column (L-column micro, CERI, Tokyo, Japan), and fractionated on an in-house fabricated 20 cm column packed with 2 µm octadecyl silane particles (CERI). Elution was performed with a linear gradient of 5% to 35% solvent B over 70 min at a flow rate of 200 nl/min (solvent A = 0.1% formic acid; solvent B = 0.1% formic acid in acetonitrile) with the use of a Dionex Ultimate 3000 HPLC System (Thermo Fisher Scientific). Eluted peptides were sprayed with a nano-electrospray source and with a column oven set at 42°C (AMR, Tokyo, Japan). The Q Exactive Hybrid Quadrupole-Orbitrap mass spectrometer plus was operated in data-independent acquisition mode. All data were acquired in profile mode using positive polarity. MS1 spectra were collected in the range of 430–860 m/z at a resolution of 35,000 using an Automated Gain Control (AGC) target value of 1 × 10^6^ ions with a maximum injection time for 50 msec. MS2 spectra were collected in the range of > 200 m/z at a resolution of 17,500 using an AGC target value of 1 × 10^6^ ions and the automatic maximum ion injection times. Twenty-one DIA windows of 20 units were ranged from 430 to 850 m/z with an overlap of 1 unit. The normalized collision energy was set to 25. All DIA raw data was processed with DIA-NN (ver. 1.8)^50^ using library-free search mode against human UniProt sequences (https://www.uniprot.org/; accessed 27 April 2021). Enzyme specificity was set to “trypsin”. Precursor m/z range was set from m/z 430 to 850. Carbamidomethylation of cysteine was set as a fixed modification.

### Animals

Male C57BL/6J mice (Jackson Laboratory, Bar Harbor, ME, USA) were used in this study. All the procedures were performed in accordance with protocols approved by the Institutional Animal Care and Use Committee of Niigata University (#SA00656) and ARRIVE guidelines. Mice were housed in groups before surgery and singly after surgery under standard laboratory conditions (21 ± 2 °C; 50% ± 10% humidity; 12-hr light/dark cycle, with darkness from 8 p.m. to 8 a.m.). Pelleted food and sterilized tap water were provided ad libitum.

### DNA damage induction in mice

Male mice received intraperitoneal injections of 200 mg.kg^-1^ etoposide (CAS no. 33419-42-0, Tokyo Chemical Industry) dissolved in 5% DMSO/PBS. Control mice received similar amounts of 5% DMSO/PBS only. After etoposide injection, moistened food pellets were placed at the bottom of all cages, and precautions were taken to minimize animal suffering during the procedures. For experiments involving APP knockdown and APP overexpression, mice were injected with AAV, followed by the administration of etoposide two weeks later.

### AAV constructs and production

pAAV plasmids were generated as follows. For pAAV-hSyn-mCherry, mCherry of pmCherry-C1 plasmid (Clontech, Mountain View, CA, USA) was subcloned into pAAV-hSyn-EGFP (#50465, Addgene) in substitution of EGFP. For APP knockdown experiments, hSyn promoter of pAAV-hSyn-mCherry was subcloned into pAAV-CMV-EGFP-U6-Control_shRNA (pAAV(shRNA)-EGFP-U6>Scramble_shRNA; Vector builder, Chicago, IL, USA) or pAAV-CMV-EGFP-U6-m*App*_shRNA (pAAV(shRNA)-EGFP-U6>m*App*_shRNA; Vector builder) in substitution of CMV with a Klenow reaction, to obtain pAAV-hSyn-EGFP-U6-Control_shRNA (Control KD) and pAAV-hSyn-EGFP-U6-m*App*_shRNA (APP KD), where the hSyn promoter drives EGFP expression in neurons. For APP overexpression experiments, APP 695 and APP Swe/Ind were amplified from pCAX APP 695 (#30137, Addgene) and pCAX APP Swe/Ind (#30145, Addgene), respectively, with primers designed with the Kozak sequence at the 5’ and a hemagglutinin (HA) tag at the 3’ terminal (forward primer sequence: *GGGTGATCAGCCACCATGGTACCCCTACGACGTGCCCGACTACGCCCTGCCCGGTTTGGCACTGCT*; reverse primer sequence: *CCCCAAGCTTCTAGTTCTGCATCTGCTCAAAGA*). They were then subcloned into pAAV-hSyn-mCherry in substitution of mCherry.

Human embryonic kidney 293T cells (#632273, Takara Bio) were used to produce AAVs. The cell identity was confirmed by short tandem repeat profiling, and the cells were tested negative for mycoplasma contamination. Cells were maintained in DMEM supplemented with 10% v/v fetal bovine serum at 37°C and 5% CO_2_. They were transfected with pAAV, pAAV2/1 (Penn Vector Core, Philadelphia, PA, USA) and pHelper plasmids (Takara Bio)^52^. AAVs in the cells and supernatant were purified by iodixanol gradient and were concentrated in phosphate-buffered saline (PBS) containing 0.001% Pluronic F68 using Amicon Ultra 100 K filter units (Merck). The titer of AAV was determined with a real-time PCR thermal cycler (Dice, Takara Bio). The resultant AAV1-Syn-mCherry (2.8 × 10^12^ GC/ml), AAV1-Syn-EGFP-Control-shRNA (8.5 × 10^12^ GC/ml), AAV1-Syn-EGFP-m*App*-shRNA (7.3 × 10^12^ GC/ml), AAV-Syn-APP 695-HA (2.6 × 10^12^ GC/ml) and AAV-Syn-APP 695Swe/Ind-HA (4.1 × 10^12^ GC/ml) were stored at −80°C until use. The expression or reduction of the target proteins was tested using immunohistochemistry.

### AAV injection into mice

Injections were performed as previously reported^52–54^ with modifications. Mice at 10 weeks were anesthetized with the anesthetic mixture of medetomidine (Nippon Zenyaku Kogyo, Koriyama, Japan), midazolam (Astellas Pharma, Tokyo, Japan) and butorphanol (Meiji Seika Pharma, Tokyo, Japan). Once anesthesia was confirmed by the loss of righting reflex, the scalp was incised and the animal placed in a stereotaxic frame (Narishige SR-6M, micromanipulator SM-15L, Narishige, Tokyo, Japan). Next, a small hole was made on the skull at the corresponding injection site by using a 27 G needle. AAV was injected into the right sensorimotor cortex (depth of 0.5 mm; coordinates, 0 mm posterior, 1.5 mm lateral to the bregma^55^; total volume, 0.6 μl) using a Hamilton small-gauge (701RN) needle (33-gauge, small hub RN needle (Ref#7762-06), Hamilton, Reno, NV, USA). The following AAVs were used for injections: control KD, AAV1-Syn-EGFP-Control-shRNA (8.5 × 10^12^ GC/ml); APP KD, AAV1-Syn-EGFP-m*App*-shRNA (7.3 × 10^12^ GC/ml); control, AAV1-Syn-mCherry (2.8 × 10^12^ GC/ml); APP WT overexpression, AAV-Syn-APP 695-HA (2.6 × 10^12^ GC/ml) and APP Swe/Ind overexpression, AAV-Syn-APP 695Swe/Ind-HA (4.1 × 10^12^ GC/ml). After the injections, the scalp was sutured, and the mice were returned to the home cages to awake.

### Immunohistochemistry of mouse brains

The animals were transcardially perfused with 4% PFA in 0.1 M PBS. The brain was carefully dissected and postfixed in the same fixatives overnight. The tissues were then cryopreserved in 30% sucrose in PBS overnight and embedded in Tissue-Tek OCT compound (Sakura Finetek Japan, Tokyo, Japan). Serial coronal 10-μm-thick sections were made with a cryostat (Leica Microsystems) and mounted on MAS-coated slides (Matsunami Glass, Kishiwada, Japan). Sections were briefly washed and were incubated in 10 mM sodium citrate buffer (pH 6.0) at 121°C for 15 min in an autoclave (HA-300MIV, HIRAYAMA). After the samples were allowed to cool gradually to RT, the sections were washed three times with PBS for 5 min each and incubated with 2% (w/v) BSA in PBS for 30 min. The sections were incubated with primary antibodies in 2% (w/v) BSA in PBS overnight at 4°C. After 3 washes with PBS, the sections were incubated with corresponding secondary antibodies diluted in blocking buffer for 2 hrs at room temperature. Samples were washed and counterstained with Hoechst 33258 (Dojindo) to stain nuclei and coverslipped with ProLong Antifade mounting medium (Thermo Fisher Scientific) and imaged with an A1R+ confocal microscope (Nikon Solutions). Negative controls (incubated with secondary but no primary antibody) were included for every analysis. The details of the antibodies used are listed in Supplementary Table 3.

### Statistics and Reproducibility

Animal experiments were not randomized, as comparisons were made between mice of different groups (for example, control versus etoposide-treated mice). Data were not acquired with blinding to experimental condition but were subjected to strict identical analyses and compared statistically using JMP software version 15.0 (SAS Institute Japan, Tokyo, Japan) or Prism 9 (GraphPad Software, Boston, MA, USA). Two-sided Student’s t-tests were used to compare arithmetic means between two groups. Pearson’s chi-squared test was applied to sets of categorical data. One-way ANOVA and post hoc analysis using Bonferroni’s correction for multiple tests were used unless indicated otherwise (Dunnett’s test was applied in some cases). All the experiments associated with the main findings were replicated at least twice independently. For animal experiments, *n* values represent the number of mice. Representative confocal images shown in the figures were independently replicated in at least six mice in each group. Images were acquired under the same microscope settings and analysed using ImageJ^56^ or Fiji software^57^. No statistical methods were used to predetermine sample size. Data are reported in the text and figures as the means ± standard errors of the means (s.e.m.). *P*-values < 0.05 were considered statistically significant. Exact *P*-values are provided in the text and figures, or statistical significance in figures is presented as \**P* < 0.05, \*\**P* < 0.01 and \*\*\**P* < 0.001. Details of statistical tests are provided in each figure and/or figure legend.

## Acknowledgements

This work was supported by the Takeda Science Foundation (Hideaki Matsui); JSPS KAKENHI Grant Numbers JP 23790744 (Hideaki Matsui), JP 22484842 (Hideaki Matsui) and JP 18955907 (Hideaki Matsui); AMED [Grant Numbers JP23gm1710010, JP22gm6110028 (Hideaki Matsui) and JP20dm0107154 (Hideaki Matsui)]; Tokyo Biochemical Research Foundation (Hideaki Matsui); Chugai Foundation for Innovative Drug Discovery Science (Hideaki Matsui); the Uehara Memorial Foundation (Hideaki Matsui) and JST [Moonshot R&D][Grant Number JPMJMS2024] (Hideaki Matsui).

## Author Contributions

G.D. performed the mice experiments, analysed the mice data, and wrote the manuscript. T.O., A.H., and M.M. performed DIA proteome analysis. Y.N. and M.U. produced the AAV vectors and provided technical assistance for stereotaxis injection. A.S. performed immunoprecipitation analysis. T.Y. and H.I. offered technical support for the mice experiments. N.M. performed cell experiments. A.N. and A.K. provided and analysed human postmortem brain samples. A.I. performed cell and related immunoblots experiments. H.M. formulated the hypothesis, acquired the grants, supervised the entire study, designed the experiments, performed the cellular experiments, analysed and interpreted the data, and wrote the manuscript.

---

**Extended Data Fig. 1.**
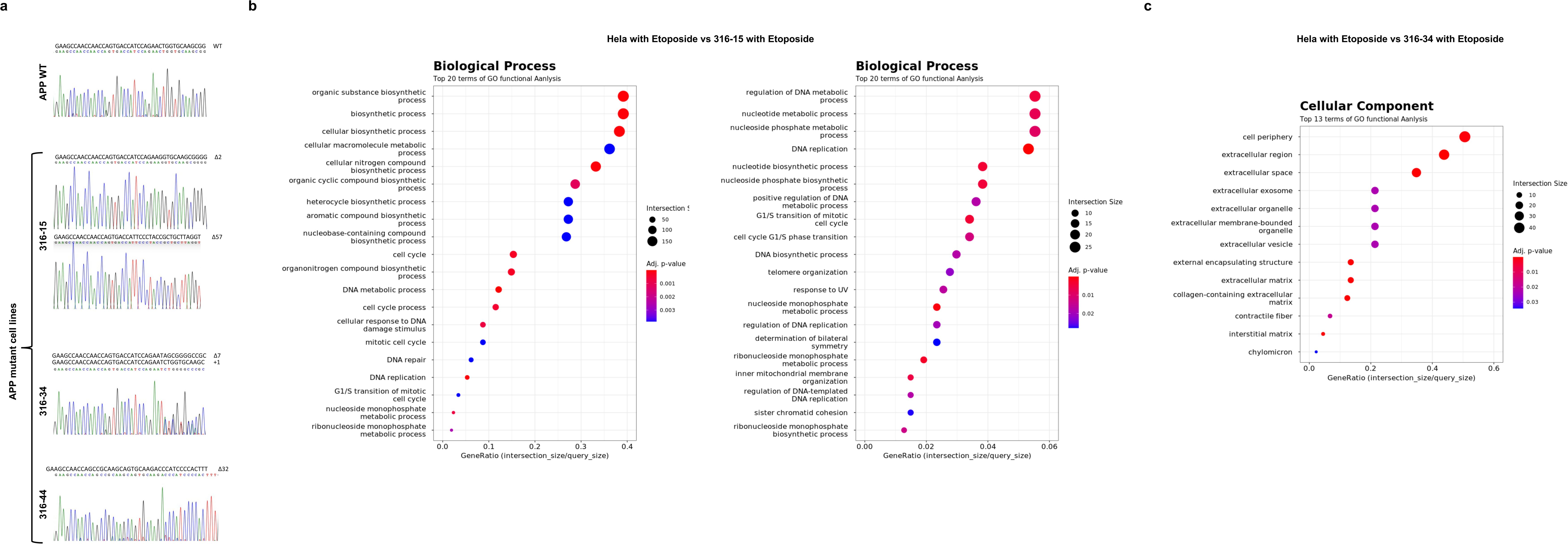
Sequence data and transcriptome of APP mutant cells. **a**, Sequence data of APP mutant cell lines (316-15, 316-34, and 316-44). 316-34 cell line is a compound heterozygote with a 7-base deletion and a 1-base insertion. 316-44 cell line is a homozygous mutation with a 32-base deletion. 316-15 cell line is compound heterozygous for 2-base deletion and 57-base deletion, with the 57-base deletion not causing a frameshift and producing a partially deleted APP protein. **b**, Gene ontology analysis of up-regulated genes in RNA sequencing of etoposide-treated 316-15 cells (20 μM, day 2) compared to etoposide-treated Hela cells. The right panel shows the size-filtered version. RNA sequencing data are attached as Source Data Extended Fig. 1b. **c**, Gene ontology analysis of down-regulated genes in RNA sequencing of etoposide-treated 316-34 cells (20 μM, day 2) compared to etoposide-treated Hela cells.

**Extended Data Fig. 2. Continued Fig. 2.**
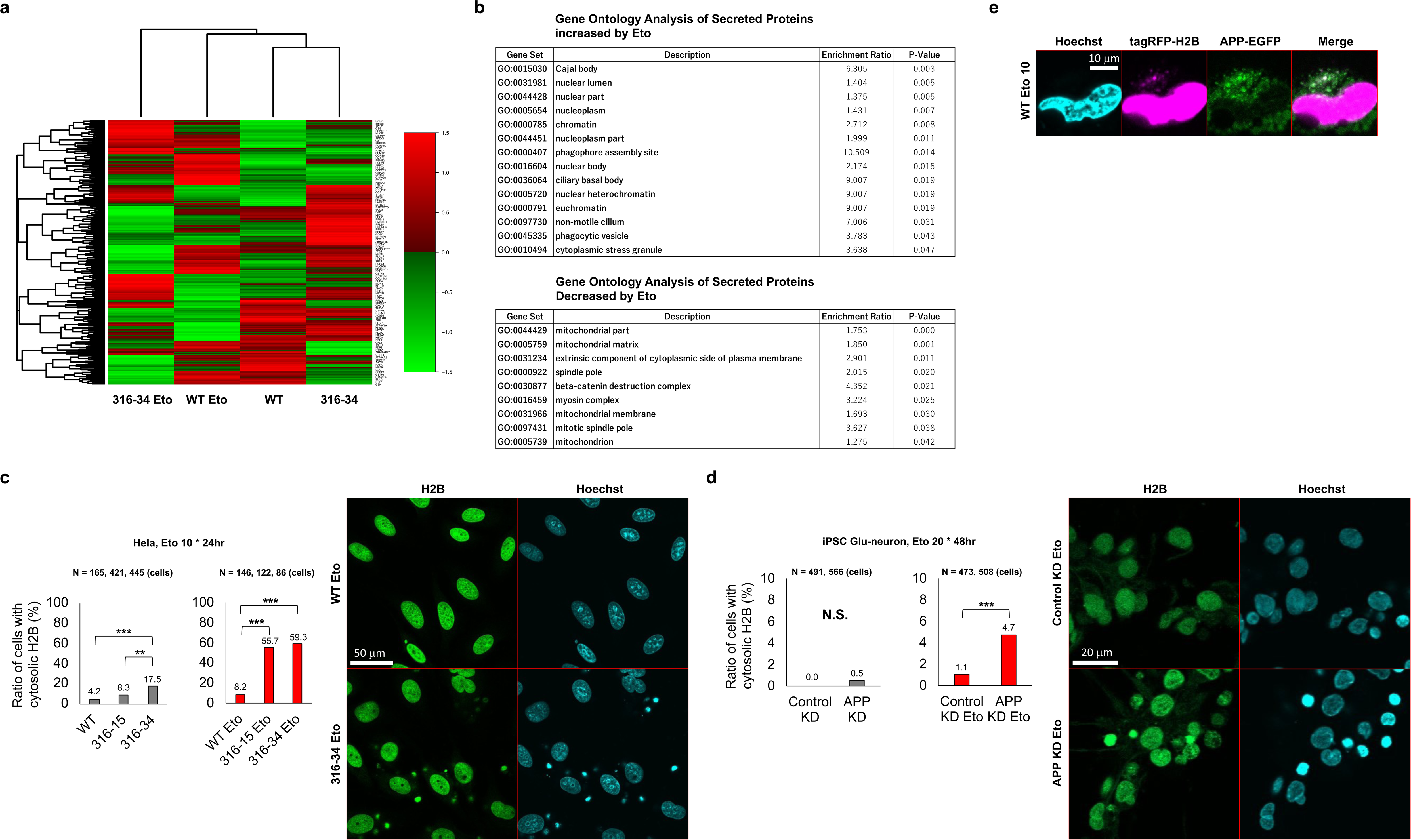
**a**, Heatmap analysis of proteins in cell culture supernatant comparing Hela and APP mutant cells (316-34) upon etoposide treatment (20 μM, day 2). **b**, Gene ontology analysis of increased (> 400%) or decreased (< 25%) proteins in cell culture supernatant of etoposide-treated Hela cells (5 μM, 24 hrs) compared to vehicle-treated Hela cells. **c**, Confocal microscopy analysis showing increased nuclear waste in etoposide-treated APP mutant cells (10 μM, 24 hrs). Statistical analysis: Pearson’s chi-squared test with Bonferroni posthoc test for multiple comparisons. **d**, Confocal microscopy analysis showing increased nuclear wastes within neurons differentiated from human iPSCs with APP knockdown and treated with etoposide (20 μM, day 2). Statistical analysis: Pearson’s chi-squared test. **e**, Snapshot of the live imaging of APP-mediated intracellular trafficking and extracellular disposal of nuclear waste. See also Supplementary Movies 3-6. N.S. non-significant, \*\**P* < 0.01 and \*\*\**P* < 0.001.

**Extended Data Fig. 3.**
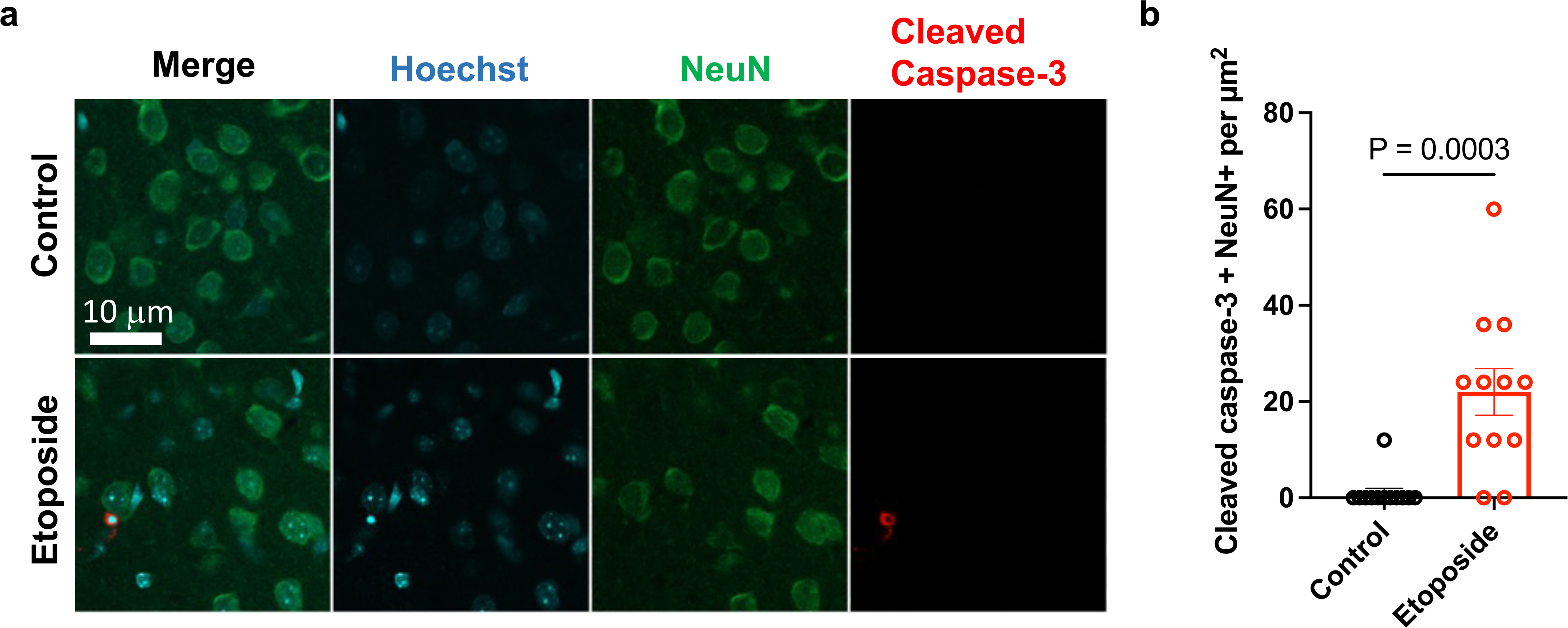
Increased neuronal death in etoposide-treated mice. **a**, Representative images of cell death immunostainings using Hoechst (nuclear marker, blue), NeuN (neuronal marker, green), and cleaved caspase-3 (apoptotic marker, red) in etoposide-treated mice and respective controls. **b**, Quantification of cleaved caspase-3 between control and etoposide-treated mice. Bars represent means; dots represent measurements for replicates (n = 6 mice per group, with 2 quantified images for each mouse). Statistical analysis: two-tailed, unpaired Student’s t-test (*P*-value is given in graph). Data are mean ± s.e.m. Values are averaged to the control group.

**Extended Data Fig. 4.**
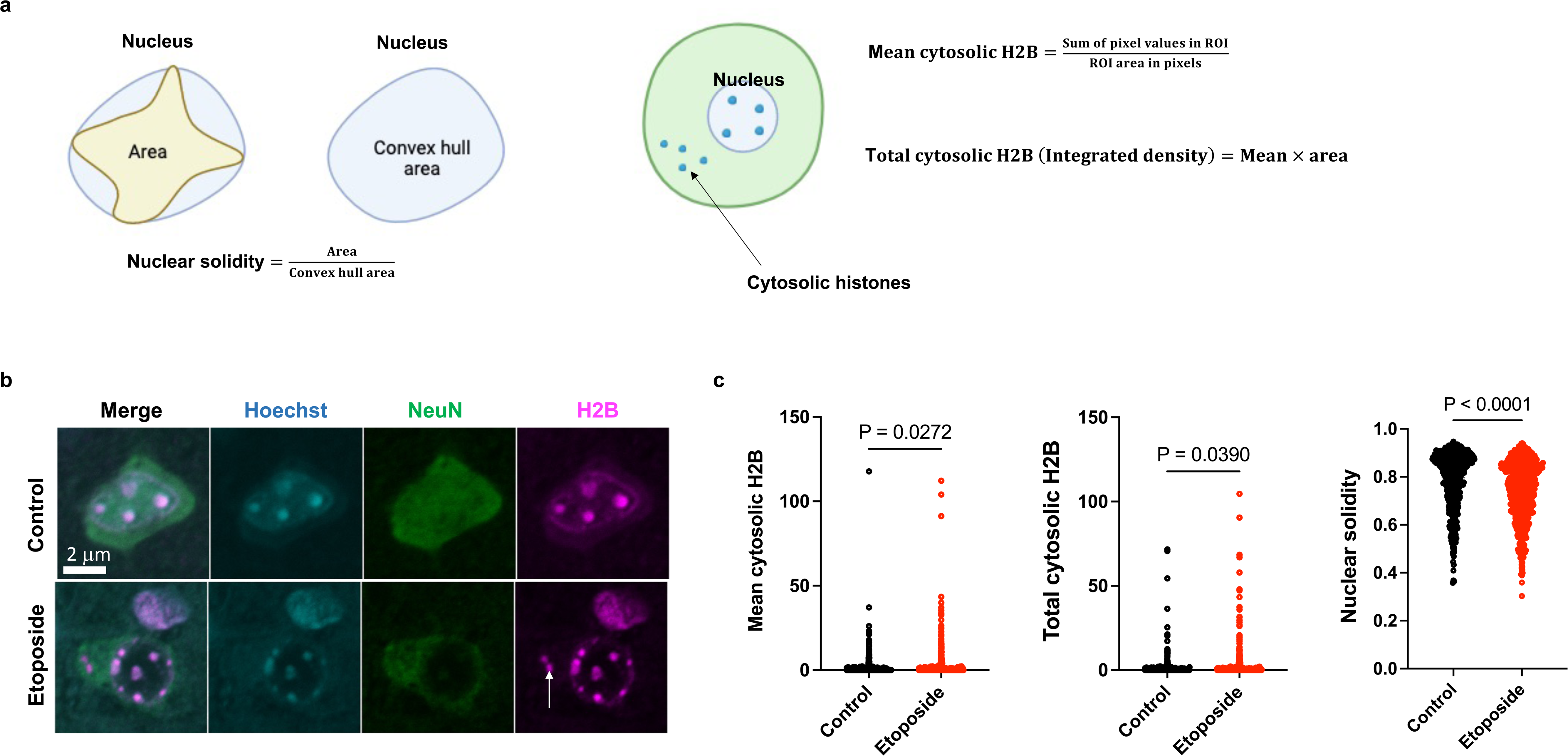
Cytosolic histone H2B in neurons of mouse brains following etoposide treatment. **a**, Schematic representation of the parameters measured relative to histones. Regions of interest (ROI) were drawn around the cytoplasm (cell excluding the nuclear compartment) and mean cytosolic H2B (ratio of the sum of pixel values by the area in pixels), total cytosolic H2B (the integrated density) and nuclear solidity (the ratio of the measured area of the nucleus to the area of its convex hull shape) were computed using Fiji; created with BioRender. **b**, Representative images of cytosolic histones immunostaining using Hoechst (nuclear marker, blue), NeuN (neuronal marker, green), and histone H2B (histone marker, magenta) in etoposide-treated mice and respective controls. White arrow indicates cytosolic H2B. **c**, Quantification of mean cytosolic H2B (left), total cytosolic H2B (middle), and nuclear solidity (right) between control and etoposide-treated mice. Violin plots show the distributions for biological replicates (n = 6 mice per group, with 2 quantified images for each mouse). Statistical analysis: two-tailed, unpaired Student’s t-test (*P*-values are given in graphs). Data are mean ± s.e.m. Values are averaged to the control group.

**Extended Data Fig. 5.**
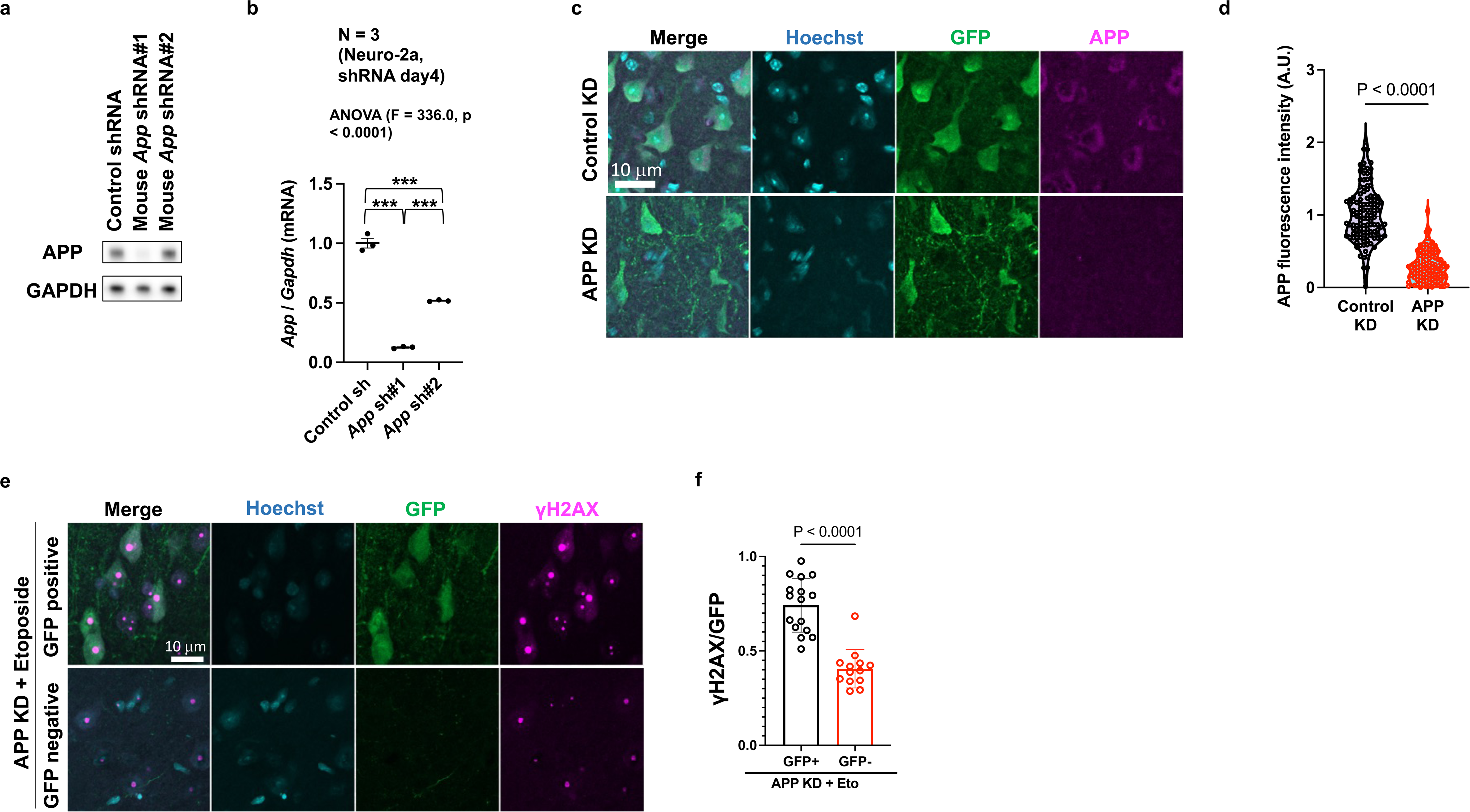
Decreased APP levels in neurons upon successful knockdown in mouse brains and comparison of DNA damage between AAV-injected/non-injected brain regions. **a**, Western blotting analysis of mouse APP under shRNA transfection into mouse Neuro-2a cells. Uncropped immunoblot is provided in Supplemental Fig. 5a. **b**, RT-qPCR analysis of mouse *App* levels under shRNA transfection into mouse Neuro-2a cells, relative to *Gapdh*. **c**, Representative immunofluorescence images of mouse cortex using Hoechst (nuclear marker, blue), GFP (neuronal marker, green), and APP (magenta) in control KD and APP KD groups. **d**, Quantification of APP mean fluorescence intensity between control KD and APP KD groups. Violin plots show the distributions for biological replicates (n = 6 mice per group, with 2 quantified images for each mouse). Statistical analysis: two-tailed, unpaired Student’s t-test (*P*-value is given in graph). **e**, Representative immunofluorescence images of mouse cortex showing DNA damage in etoposide-treated APP KD mice. GFP positive region is compared with the contralateral (not injected) side showing increased DNA damage in the ipsilateral region. **f**, Quantification of γH2AX foci between GFP positive and contralateral regions of etoposide-treated APP KD mice. Bars represent means; dots represent measurements for replicates (n = 7 mice per group, with 2-3 quantified images for each mouse). Statistical analysis: two-tailed, unpaired Student’s t-test (*P*-value is given in graph). Data are mean ± s.e.m. Values are averaged to the control KD group.

**Extended Data Fig. 6.**
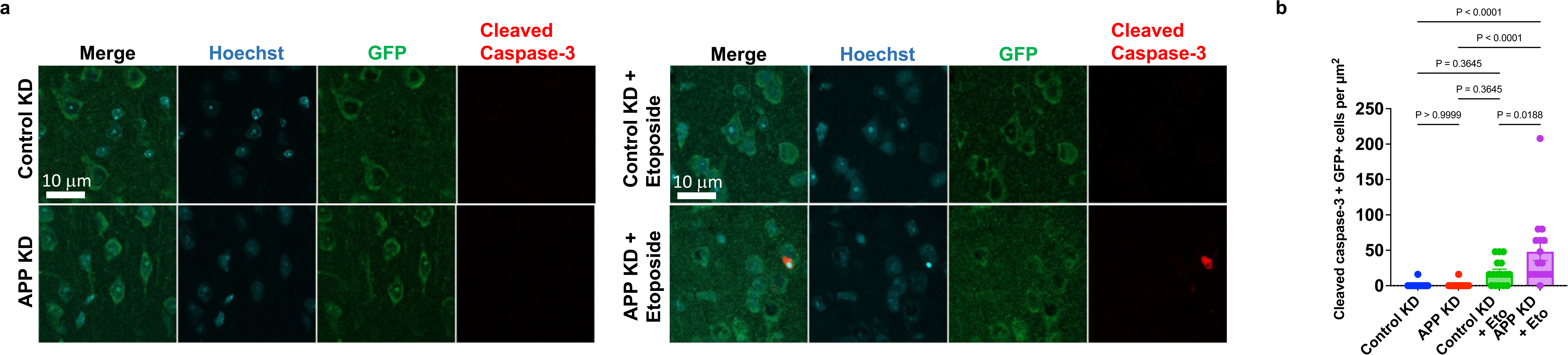
Increased neuronal death in etoposide-treated APP KD mice. **a**, Representative images of cell death immunostaining using Hoechst (nuclear marker, blue), GFP (neuronal marker, green), and cleaved caspase-3 (apoptotic marker, red) in etoposide-treated control KD, APP KD mice and their respective controls. **b**, Quantification of cleaved caspase-3 between etoposide-treated control KD, APP KD mice and their respective controls. Bars represent means; dots represent measurements for replicates (n = 8 mice per group, with 2 quantified images for each mouse). Statistical analysis: ordinary one-way ANOVA with Bonferroni posthoc test for multiple comparisons (*P*-values for the posthoc tests are given in the graphs). Data are mean ± s.e.m. Values are averaged to the untreated control KD group.

**Extended Data Fig. 7.**
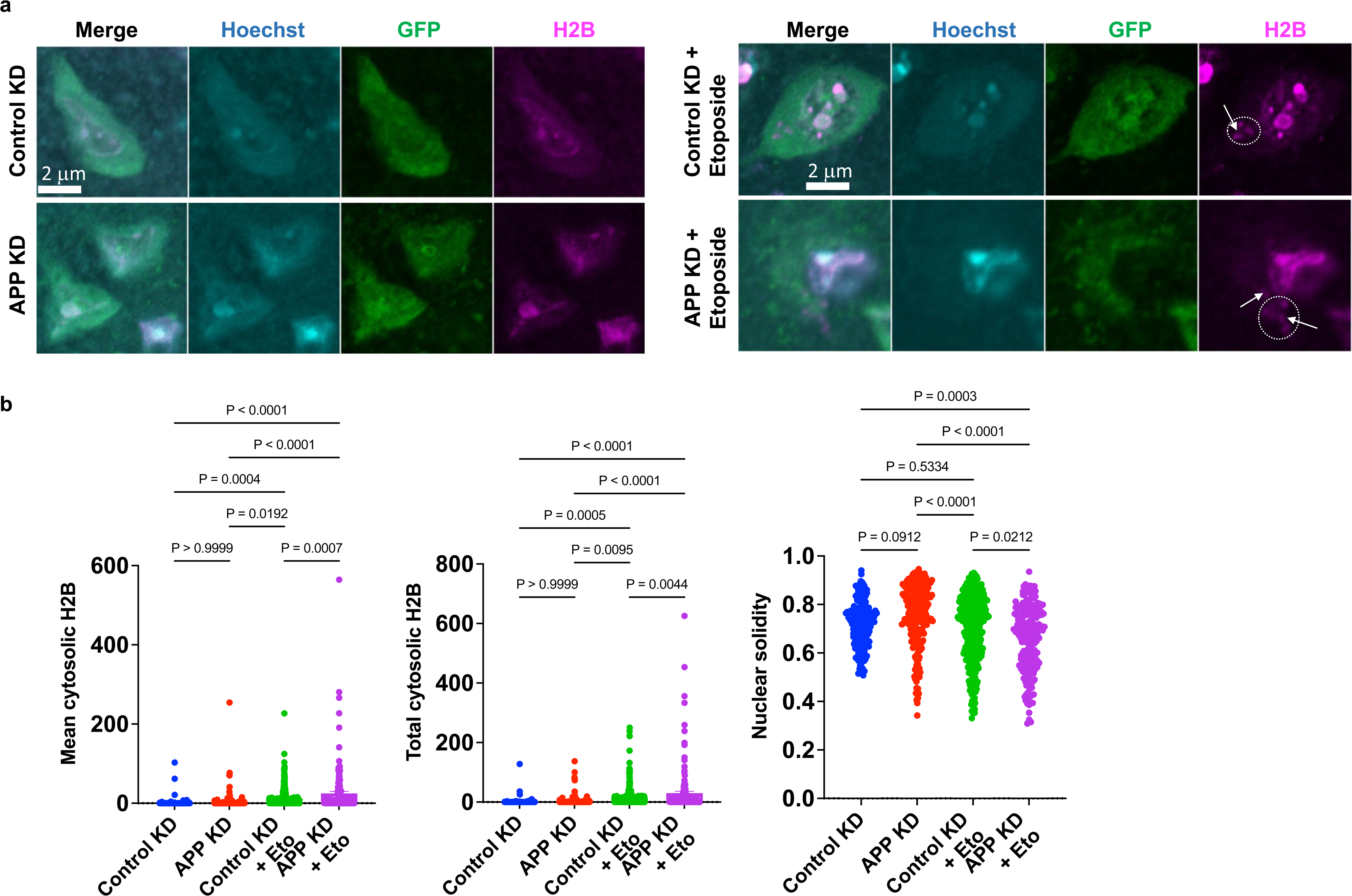
Increased cytoplasmic histone H2B in neurons of mouse brains following APP KD and etoposide treatment. **a**, Representative images of cytosolic histones immunostaining using Hoechst (nuclear marker, blue), GFP (neuronal marker, green), and histone H2B (histone marker, magenta) in etoposide-treated control KD, APP KD mice and their respective controls. White arrows indicate cytosolic H2B. **b**, Quantification of mean cytosolic H2B (left), total cytosolic H2B (middle) and nuclear solidity (right) between etoposide-treated control KD, APP KD mice and their respective controls. Violin plots show the distributions for biological replicates (n = 6 mice per group, with 2 quantified images for each mouse). Statistical analysis: ordinary one-way ANOVA with Bonferroni posthoc test for multiple comparisons (*P*-values for the posthoc tests are given in the graphs). Values are averaged to the untreated control KD group. Data are mean ± s.e.m.

**Extended Data Fig. 8.**
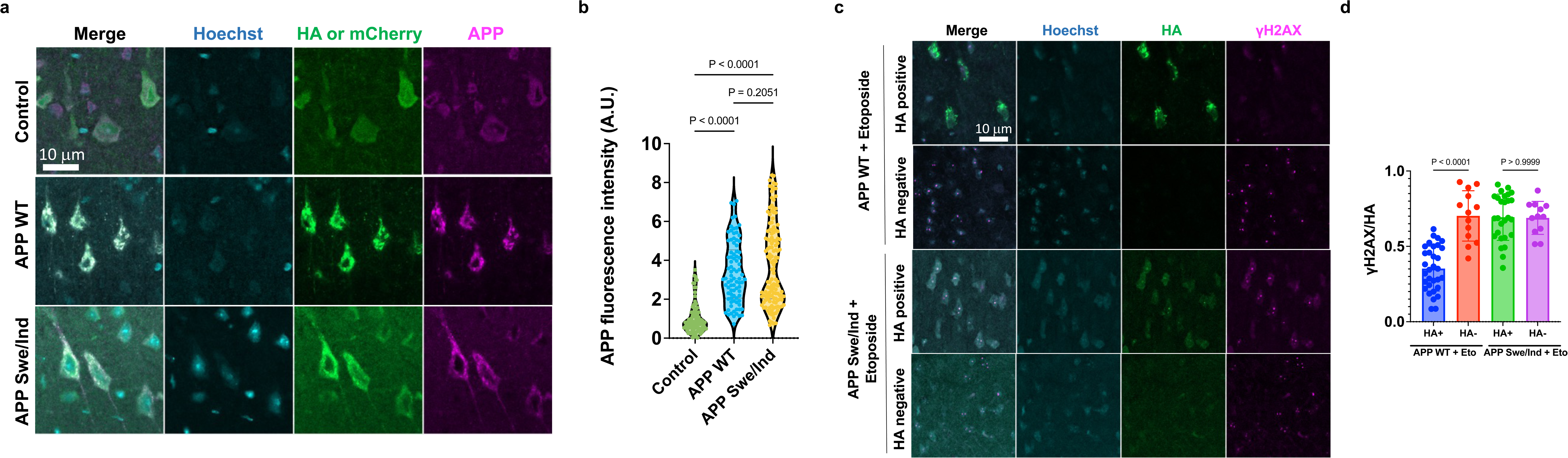
Increased APP levels in neurons following APP WT and Swe/Ind overexpression in mouse brains and comparison of DNA damage between AAV-injected/non-injected mouse brain regions. **a**, Representative immunofluorescence images of mouse cortex using Hoechst (nuclear marker, blue), HA or mCherry (neuronal marker, green), and APP (magenta) in control, APP WT, and APP Swe/Ind groups. **b**, Quantification of APP mean fluorescence intensity between control, APP WT, and APP Swe/Ind groups. Violin plots show the distributions for biological replicates (n = 6 mice per group, with 2 quantified images for each mouse). Statistical analysis: ordinary one-way ANOVA with Bonferroni posthoc test for multiple comparisons (*P*-values for the posthoc tests are given in the graphs). **c**, Representative immunofluorescence images of mouse cortex showing DNA damage in etoposide-treated APP WT and APP Swe/Ind mice. HA positive regions are compared with the negative regions, showing increased DNA damage in the ipsilateral region for etoposide-treated APP Swe/Ind and reduced DNA damage for etoposide-treated APP WT. **d**, Quantification of γH2AX foci between HA positive and negative regions of etoposide-treated APP WT and APP Swe/Ind mice. Bars represent means; dots represent measurements for replicates (n = 7 mice per group, with 2-5 quantified images for each mouse). Statistical analysis: ordinary one-way ANOVA with Bonferroni posthoc test for multiple comparisons (*P*-values for the posthoc tests are given in the graphs). The exact *P*-values of comparison are presented in the figures, respectively. Data are mean ± s.e.m. Values are averaged to the control group.

**Extended Data Fig. 9.**
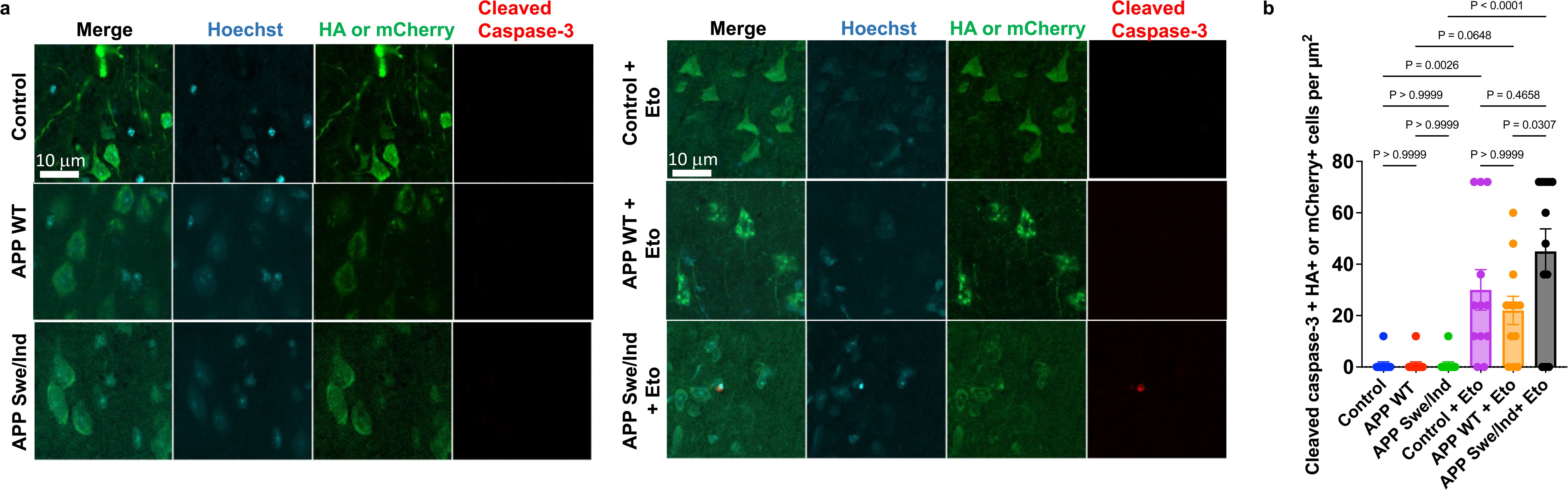
Protective effect of APP WT against etoposide-induced neuronal death in mouse brains. **a**, Representative images of cell death immunostaining using Hoechst (nuclear marker, blue), HA or mCherry (neuronal marker, green), and cleaved caspase-3 (apoptotic marker, red) in etoposide-treated control, APP WT and APP Swe/Ind mice and their respective controls. **b**, Quantification of cleaved caspase-3 between etoposide-treated control, APP WT and APP Swe/Ind mice and their respective controls. Bars represent means; dots represent measurements for replicates (n = 6 mice per group, with 2 quantified images for each mouse). Statistical analysis: ordinary one-way ANOVA with Bonferroni posthoc test for multiple comparisons (*P*-values for the posthoc tests are given in the graphs). Data are mean ± s.e.m. Values are averaged to the untreated control group.

**Extended Data Fig. 10.**
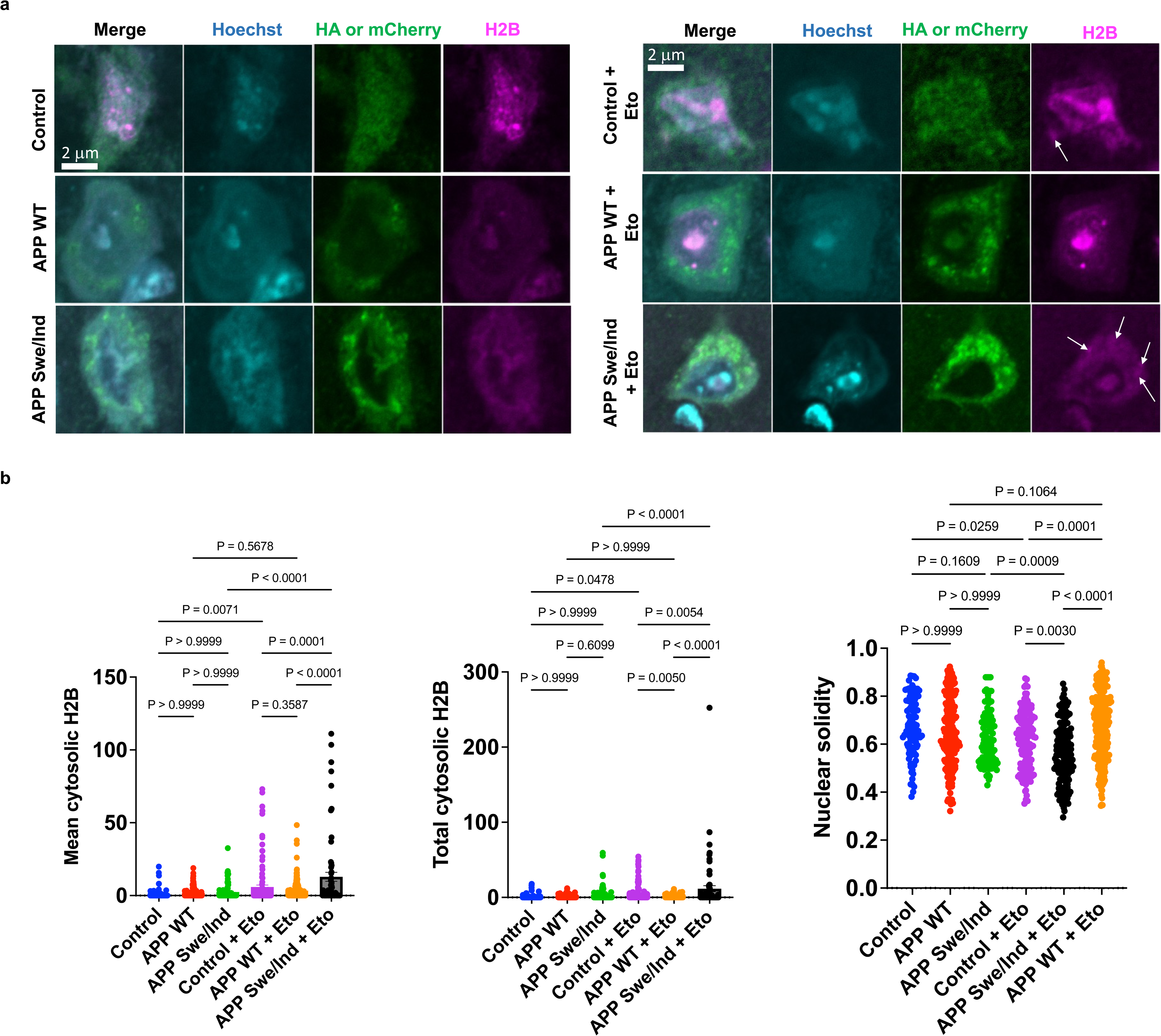
Protective effect of APP WT against etoposide-induced neuronal nuclear impairments in mouse brains. **a**, Representative images of cytosolic histones immunostaining using Hoechst (nuclear marker, blue), HA or mCherry (neuronal marker, green), and histone H2B (histone marker, magenta) in etoposide-treated control, APP WT and APP Swe/Ind mice and their respective controls. White arrows indicate cytosolic H2B. **b**, Quantification of mean cytosolic H2B (left), total cytosolic H2B (middle), and nuclear solidity (right) between etoposide-treated control, APP WT and APP Swe/Ind mice and their respective controls. Violin plots show the distributions for biological replicates (n = 6 mice per group, with 2 quantified images for each mouse). Statistical analysis: ordinary one-way ANOVA with Bonferroni posthoc test for multiple comparisons (*P*-values for the posthoc tests are given in the graphs). Data are mean ± s.e.m. Values are averaged to the untreated control group.

**Extended Data Fig. 11.**
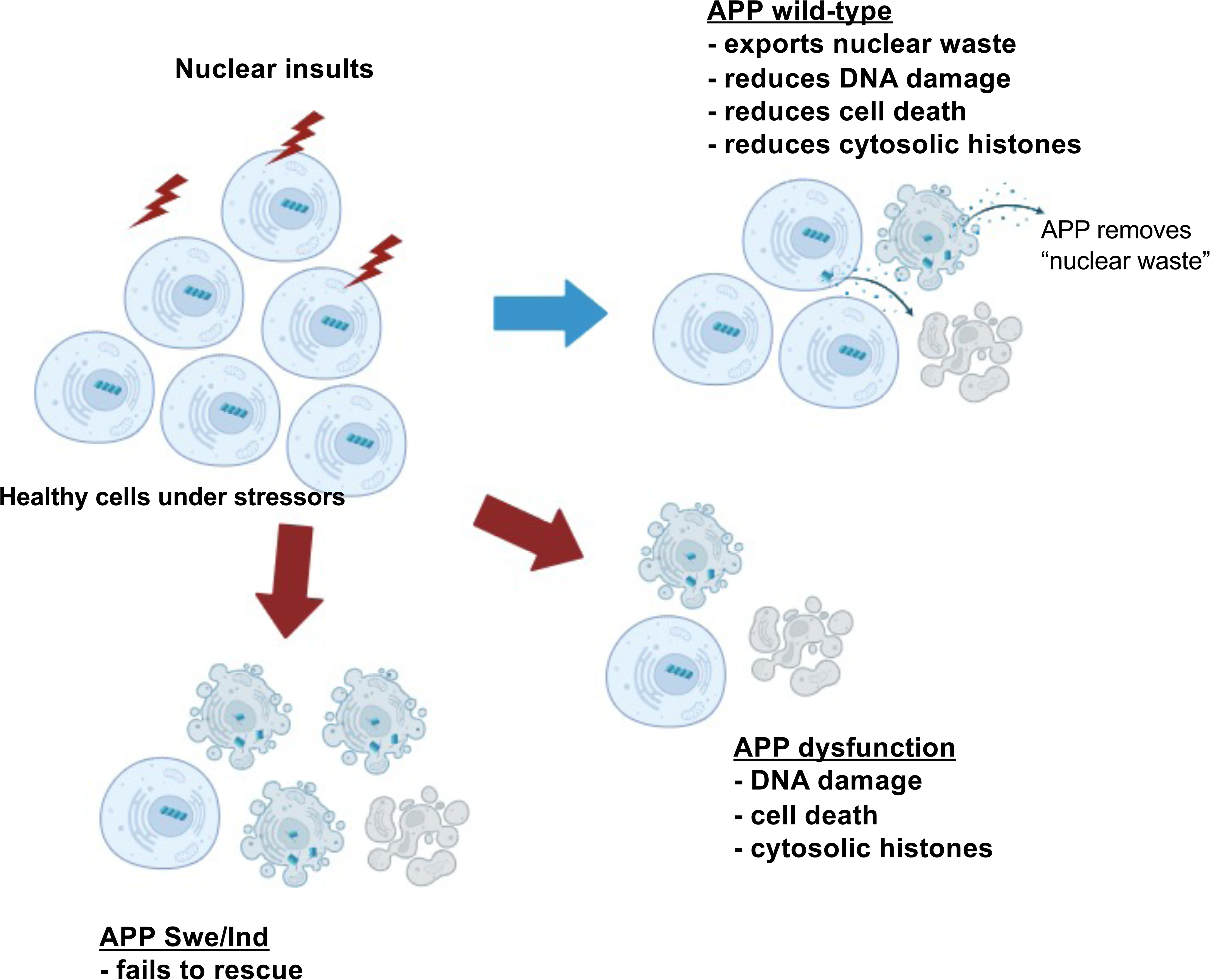
Schematic of APP protective effects in response to nuclear insults. The scheme illustrates how APP dysfunction acts as a driving force for enhanced DNA damage, cell death, and nuclear impairment; created with BioRender. It is noteworthy that APP WT, as opposed to the APP Swe/Ind mutant, ameliorates the substantial alterations induced by nuclear insults. This rescue mechanism of APP WT is due to the removal of cytoplasmic waste derived from the damaged nucleus, referred to as “nuclear waste”; a concerted action that results in a mitigation of DNA damage, a reduction in cell death, and a decrease in nuclear abnormalities.

