## Supplementary Information for "Extracellular disposal of nuclear waste by APP: a protective mechanism impaired in Alzheimer’s disease"

**Supplementary Information Guide**

**SI Figure 1.** Full western blotting images related to Fig. 1c. The area covered by the gray rectangle is the western blotting of other experiments and is not relevant to this paper. Red boxes indicate cropped regions as presented in the figures.

**SI Figure 2.**  **a**, Full western blotting images related to Fig. 2a. Red boxes indicate cropped regions as presented in the figures. **b**, Full western blotting images related to Fig. 2d. Red boxes indicate cropped regions as presented in the figures.

**SI Figure 3.** Full western blotting images related to Fig. 3d. The area covered by the gray rectangle is the western blotting of other experiments and is not relevant to this paper. Red boxes indicate cropped regions as presented in the figures.

**SI Figure 4.** **a,** Full western blotting images related to Fig. 4a. The area covered by the gray rectangle is the western blotting of other experiments and is not relevant to this paper. Red boxes indicate cropped regions as presented in the figures. **b,** Full western blotting images related to Fig. 4b. The area covered by the gray rectangle is the western blotting of other experiments and is not relevant to this paper. Red boxes indicate cropped regions as presented in the figures.

**SI Figure 5. a,** Full western blotting images related to Extended Data Fig. 5a. The area covered by the gray rectangle is the western blotting of other experiments and is not relevant to this paper. Red boxes indicate cropped regions as presented in the figures. **b**, Full western blotting images related to Fig. 5d. Red boxes indicate cropped regions as presented in the figures.

**SI Figure 6.** Full western blotting images related to Fig. 6c. Red boxes indicate cropped regions as presented in the figures.

**Supplementary Table 1.** Information on human postmortem brain cases used in this study.

**Supplementary Table 2.** Details of siRNAs used in this study.

**Supplementary Table 3.** Details of antibodies used in this study.

**Supplementary Table 4.** Details of oligonucleotides used in this study.

**Supplementary Movie 1.** 3D reconstruction of confocal images showing Histone H2B, Fyn-Venus, and Hoechst staining in Fig. 3f (Left).

**Supplementary Movie 2.** 3D reconstruction of confocal images showing ANXA5, Fyn-Venus, and Hoechst staining in Fig. 3f (Right).

**Supplementary Movie 3.** Live imaging of APP-mediated intracellular trafficking and extracellular disposal of nuclear waste. This movie shows the dynamic movement of APP. The width is 62.37 μm.

**Supplementary Movie 4.** Live imaging of APP-mediated intracellular trafficking and extracellular disposal of nuclear waste. This movie shows the dynamic movement of histone H2B. The width is 62.37 μm.

**Supplementary Movie 5.** Live imaging of APP-mediated intracellular trafficking and extracellular disposal of nuclear waste. This movie shows the bright field image. The width is 62.37 μm.

**Supplementary Movie 6.** Live imaging of APP-mediated intracellular trafficking and extracellular disposal of nuclear waste. This movie shows the merged image of APP, histone H2B and bright field. The width is 62.37 μm.
