## Supplementary figures and images for "Extracellular disposal of nuclear waste by APP: a protective mechanism impaired in Alzheimer’s disease"

### Supplementary Fig.1

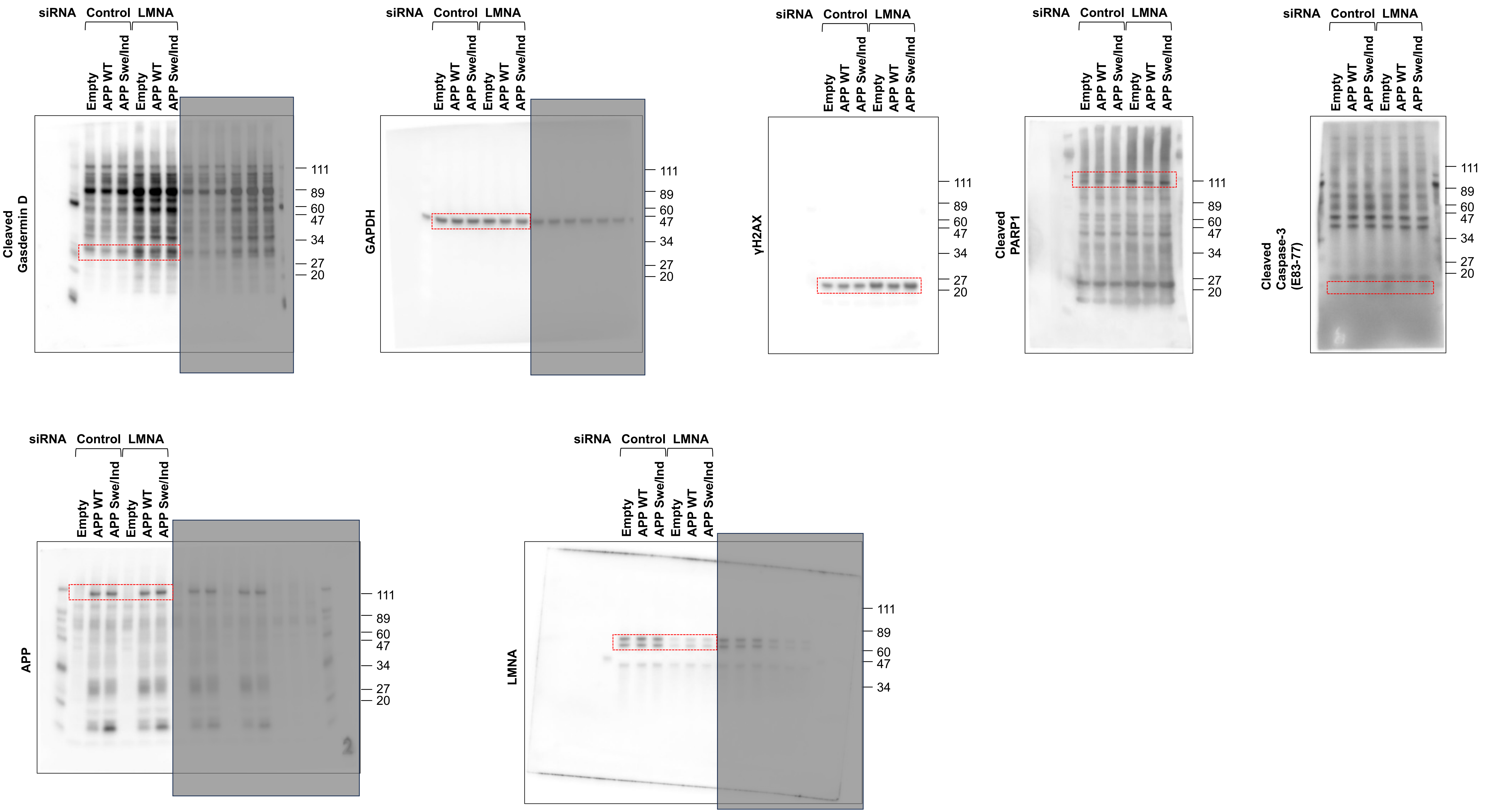

### Supplementary Fig.2

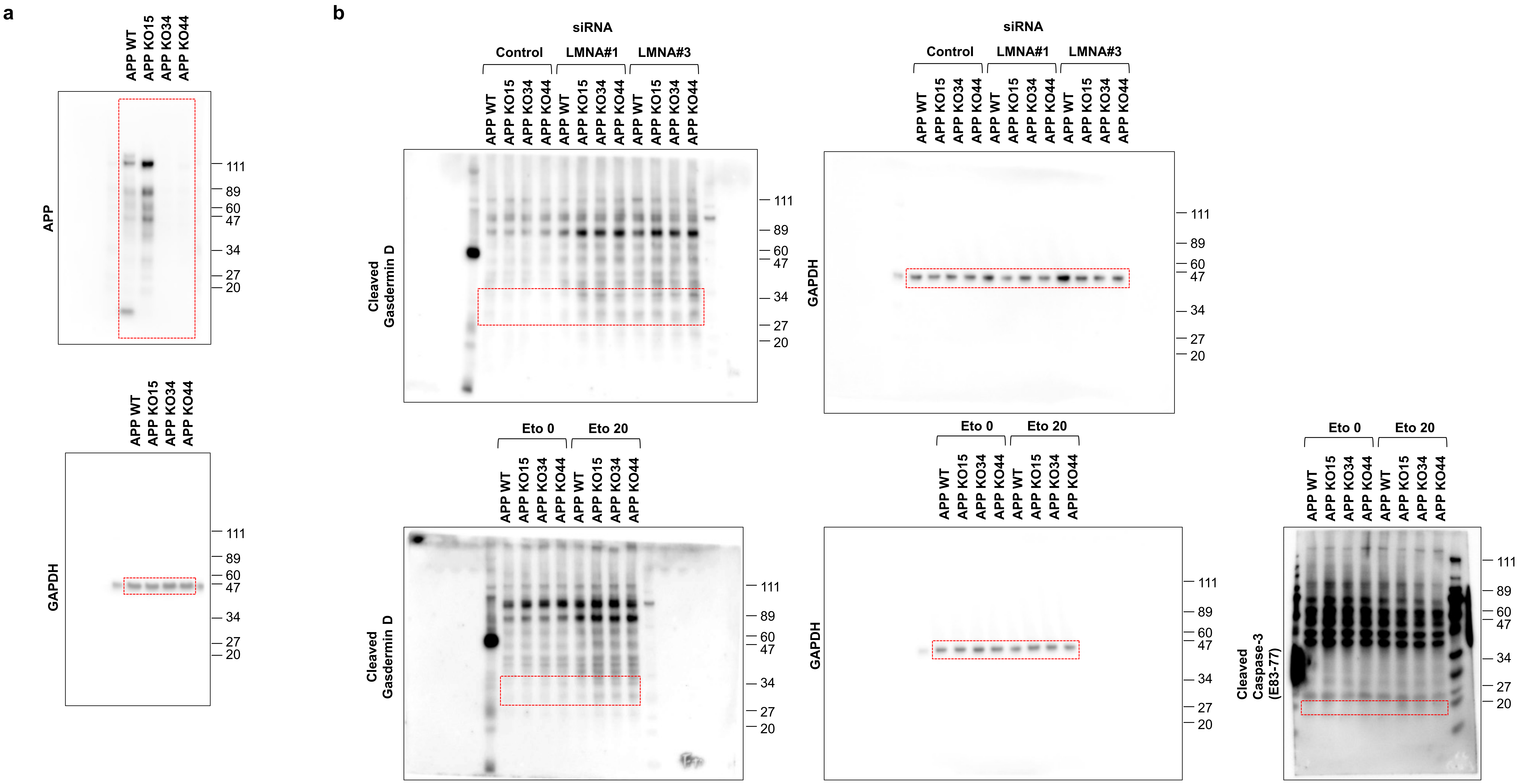

### Supplementary Fig.3

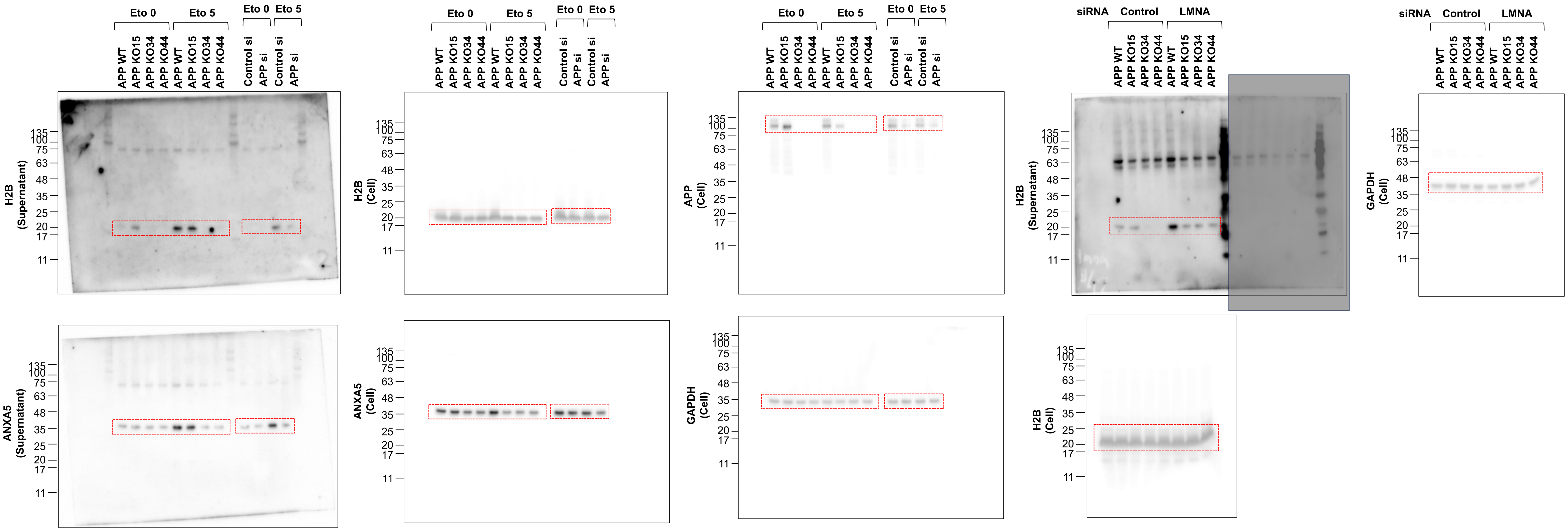

### Supplementary Fig.4

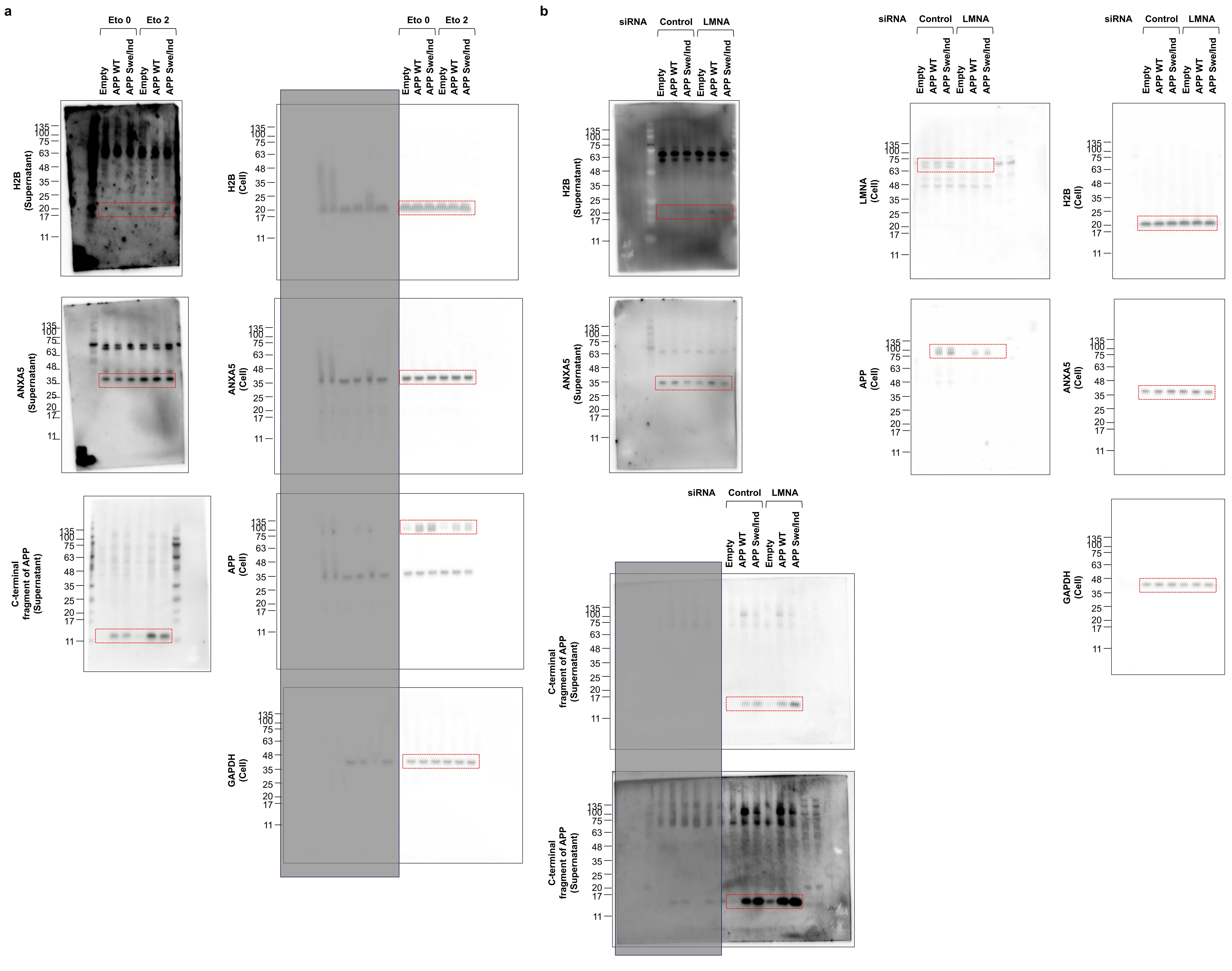

### Supplementary Fig.5

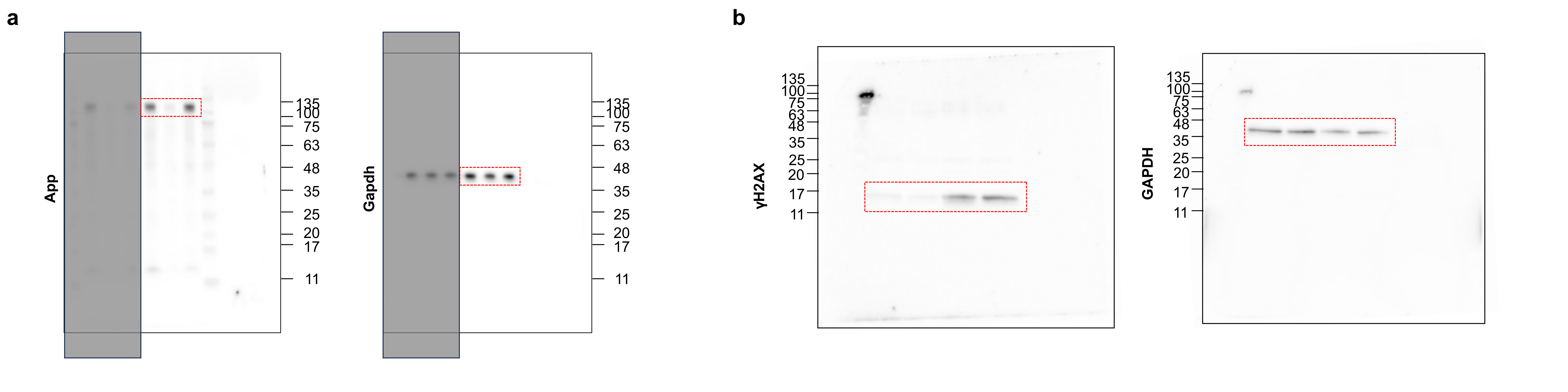

### Supplementary Fig.6

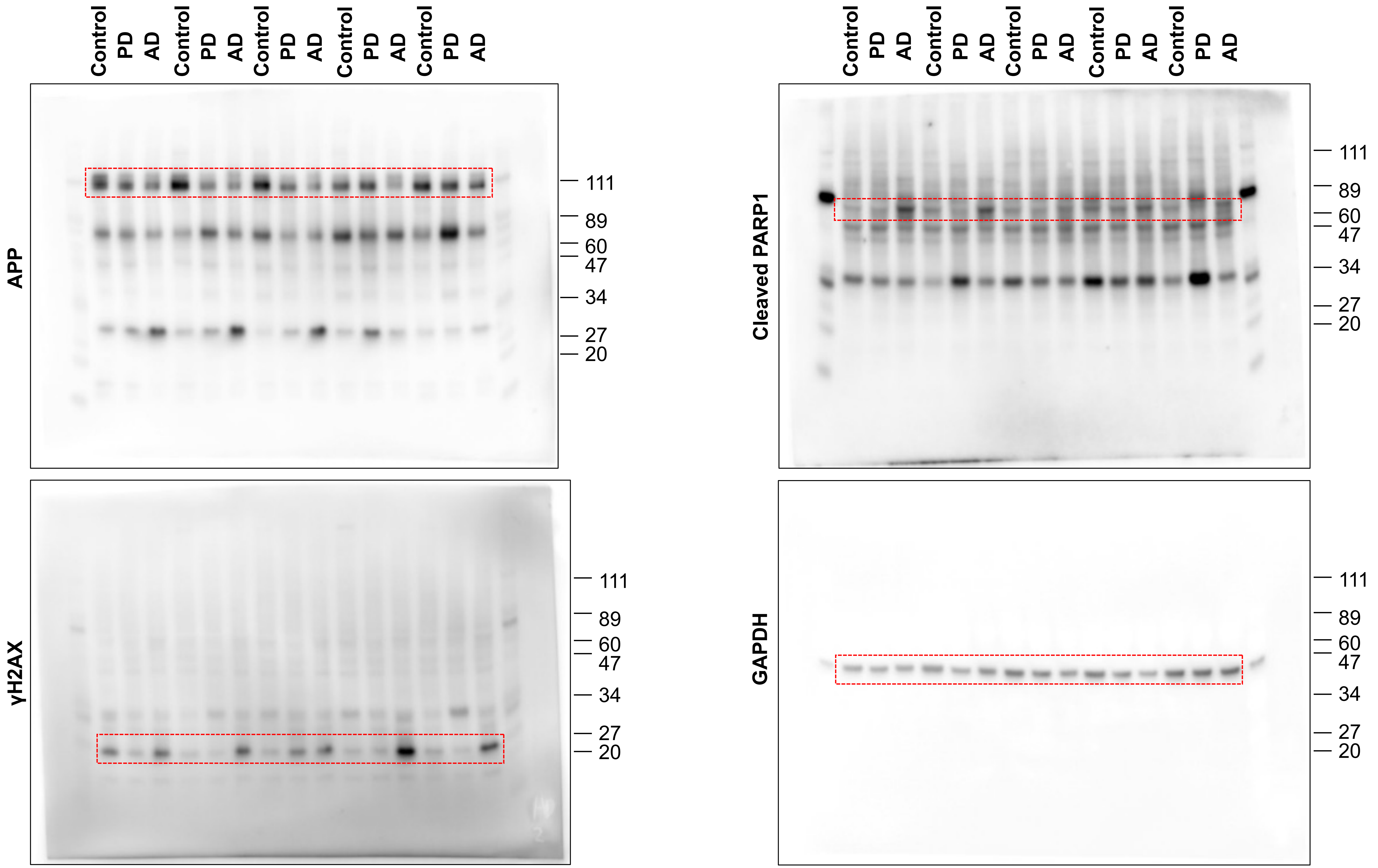
